# Blue carbon sequestration dynamics within tropical seagrass sediments: Long-term incubations for changes over climatic scales

**DOI:** 10.1101/604587

**Authors:** Chuan Chee Hoe, John Barry Gallagher, Chew Swee Theng, Norlaila Binti Mohd. Zanuri

**Affiliations:** Faculty of Science and Natural Resources, Universiti Malaysia Sabah, 88400, Sabah, Malaysia; Centre for Marine and Coastal Studies, Universiti Sains Malaysia, 11800, Penang, Malaysia; Institute for Marine and Antarctic Studies, University of Tasmania, 7000, Tasmania, Australia; Borneo Marine Research Institute, Universiti Malaysia Sabah, 88400, Sabah, Malaysia

**Keywords:** sediment geochemistry, diagenesis, carbonate, pyrogenic carbon, methane, sediment isotope tomography

## Abstract

Determination of blue carbon sequestration in seagrass sediments over climatic time scales relies on several assumptions, such as no loss of particulate organic carbon (POC) after one or two years, tight coupling between POC loss and CO_2_ emissions, no dissolution of carbonates and removal of the stable black carbon (BC) contribution. We tested these assumptions via 500-day anoxic decomposition/mineralisation experiments to capture centennial parameter decay dynamics from two sediment horizons robustly dated as 2 and 18 years old. No loss of BC was detected, and decay of POC was best described for both horizons by near-identical reactivity continuum models. The models predicted average losses of 49% and 51% after 100 years of burial and 20–22 cm horizons, respectively. However, the loss rate of POC was far greater than the release rate of CO_2_, both before and after accounting for CO_2_ from anoxic particulate inorganic carbon (PIC) production, possibly as siderite. The deficit could not be attributed to dissolved organic carbon or dark CO_2_ fixation. Instead, evidence based on δ^13^CO_2_, acidity and lack of sulphate reduction suggested methanogenesis. The results indicate the importance of centennial losses of POC and PIC precipitation and possibly methanogenesis in estimating carbon sequestration rates.

## Introduction

Seagrasses, along with mangroves, saltmarsh and seaweeds, are increasingly touted as a significant global carbon sink (McLeod *et al*. 2011). For seagrass in particular, this service is based on two separate concepts: sedimentary carbon stocks and rates of sedimentary carbon sequestration. The stock or storage service concept, in the mitigation of greenhouse gas emissions, is a scalar concept and conceived at the meadow scale. It has traditionally been estimated by potential carbon loss to mineralisation should it be disturbed over a climatic unit of time (Pendleton *et al*. 2012). The depth of such disturbance, and the extent of its effect on the carbon stock, is dependent on the type of disturbance (Siikamäki *et al*. 2013; Gallagher 2017) and independent of the time it took the carbon to accumulate. The sediment found within seagrass beds contains a sizable organic component consisting of a mix of seagrass litter, associated epiphyte and microalgal detritus, and additional inputs from adjacent land activities and fluvial deposition as well as saltmarsh and mangrove ecosystems (Kennedy *et al*. 2010). In contrast, the carbon sequestration service is a vector concept. Rates of sequestration depend on the balance between detrital production and mineralisation relative to an alternative and likely non-vegetated state (Siikamäki *et al*. 2013; Gallagher 2017). Non-vegetated sediments have in general shown increased rates of mineralisation (Kristensen *et al*. 1995) and mobilisation of dissolved organic carbon (DOC) during resuspension (Koelmans and Prevo 2003). As this is a service in the mitigation of global warming, its extent has been traditionally estimated as the rate at which sedimentary organic mass accumulates over time scales ranging from inter-decadal to centennial (Duarte *et al*. 2013, Gallagher, 2015), subsequently integrated across the meadow.

Notwithstanding uncertainties about the size of past meadow coverage and the amount and fate of exported litter (Gallagher 2014; Duarte and Krause-Jensen 2017), researchers are increasingly recognising that the traditional methods of calculating sedimentary carbon accumulation rates may have built-in biases (Gallagher 2015; Sophia and Robie 2016; Chew and Gallagher 2018). For example, previous studies have failed to subtract allochthonous recalcitrant forms of carbon such as black or pyrogenic carbon from estimated carbon stocks. Pyrogenic carbon is produced by incomplete combustion of biomass and fossil fuels. It is considered sufficiently stable to be outside the climatic carbon loop (Liang *et al*. 2008; Wang *et al*. 2016), and thus its storage and sequestration within seagrass ecosystem sediments cannot be counted as a greenhouse gas mitigation service (Chew and Gallagher 2018). Mass accumulation rates, the product of sedimentation rates and particulate organic carbon (POC) concentrations, have no assumed significant losses after one to two years within their surface sediments (Cebrian 1999). The humification of seagrass, macroalgae and mangrove detritus has been shown to occur over several months after deposition, becoming more recalcitrant after burial (Middelburg 1989; Burdige 2007). Further, any such losses are assumed to be tightly coupled with CO_2_ emissions, ostensibly from aerobic mineralisation or sulphate reduction (Burdige 1991) whereby the release of ammonia can feed further production. Methanogenesis has been known to play a measurable role within highly organic non-vegetated coastal sediments (Boehme *et al*. 1996). However, long-term incubation experiments with marine non-vegetative sediments consisting of predominantly, but not exclusively, phytoplanktonic sources suggest that POC continues to be lost within deeper and older sediments (Westrich and Berner 1984; Burdige 1991; Arndt *et al*. 2013; Canuel *et al*. 2017), with further losses of the POC fraction transformed to a mobile DOC pool (Holmer 1996; Hee *et al*. 2001; Burdige *et al*. 2016). Furthermore, the CO_2_ need not be from organic mineralisation. Sulphate reduction within non-vegetated coastal sediments has been found to result in sufficient alkalisation to produce CO_2_ from the subsequent precipitation of CaCO_3_ in the form of particulate inorganic carbon (PIC) (Mucci *et al*. 2000; Rassmann *et al*. 2016). Should this be a phenomenon within anoxic seagrass sediments, then this apparent emission source needs to be balanced with PIC dissolution subsequent to re-alkalisation of the water column after disturbance of the non-vegetated state. This can reduce the water column’s *p*CO_2_, which ironically becomes a net CO_2_ sink from the atmosphere, the extent of which depends on the residence time of the water body (Howard *et al*. 2017).

### Aims

This study aims, for the first time, to use long-term (500 days) ‘open’ anoxic slurry incubations to capture the rates and dynamics of POC and black carbon (BC) mineralisation and decomposition within highly organic seagrass meadow sediments in a tropical climate. Incubation was followed by a relatively short period of aeration (30 days) as a model for the immediate effects of disturbance on the mineralisation and decomposition of both POC and PIC. Younger (1–2 years old) surface sediments were used to compare the POC and PIC decomposition and mineralisation rates with that of deeper, older horizons. This is done by fitting the time series to the most appropriate diagenetic model (Arndt *et al*. 2013). After sediment deposition ages were determined with either an evaluated event or ^210^Pb geochronology, the model was used to extrapolate any losses over 100 years for a more considered rate of POC sequestration. The newly measured POC is then further constrained by measurements of additional diagenetic variables, namely CO_2_, coloured dissolved organic matter (CDOM) as a proxy for the DOC pool, ammonia as evidence of sulphate reduction and PIC, in the form of carbonate, to disentangle changes in CO_2_ from inorganic and organic dynamics.

## Materials and Methods

### Study site

Two similar subtidal *Enhalus* sp. seagrass meadows in separate branches of the Salut– Mengkabong estuary were chosen for the study (Fig. 1). The region can be considered as moderately urban; it is located 20 km north of a city centre (Kota Kinabalu, Sabah, Malaysia) and within the penumbra of the near-annual southwest Borneo and Sumatra haze events. These events ostensibly deposit BC into the estuary from peat fires on the southern part of the island as well as from slash-and-burn land-clearing activities (Gaveau *et al*. 2014; Chew and Gallagher 2018). The two bays are both turbid and shallow (1–3 m) and surrounded by mangrove forests with exposed intertidal mud banks. One meadow, within the Salut branch, was used to collect sediments for the slurry incubations, while the other meadow, within the Mengkabong branch, was used to constrain the Salut meadow’s geochronology. This was necessary for disentangling and identifying likely and known regional storm depositional events from unknown local disturbances (Gallagher and Ross 2018).

**Fig. 1.**
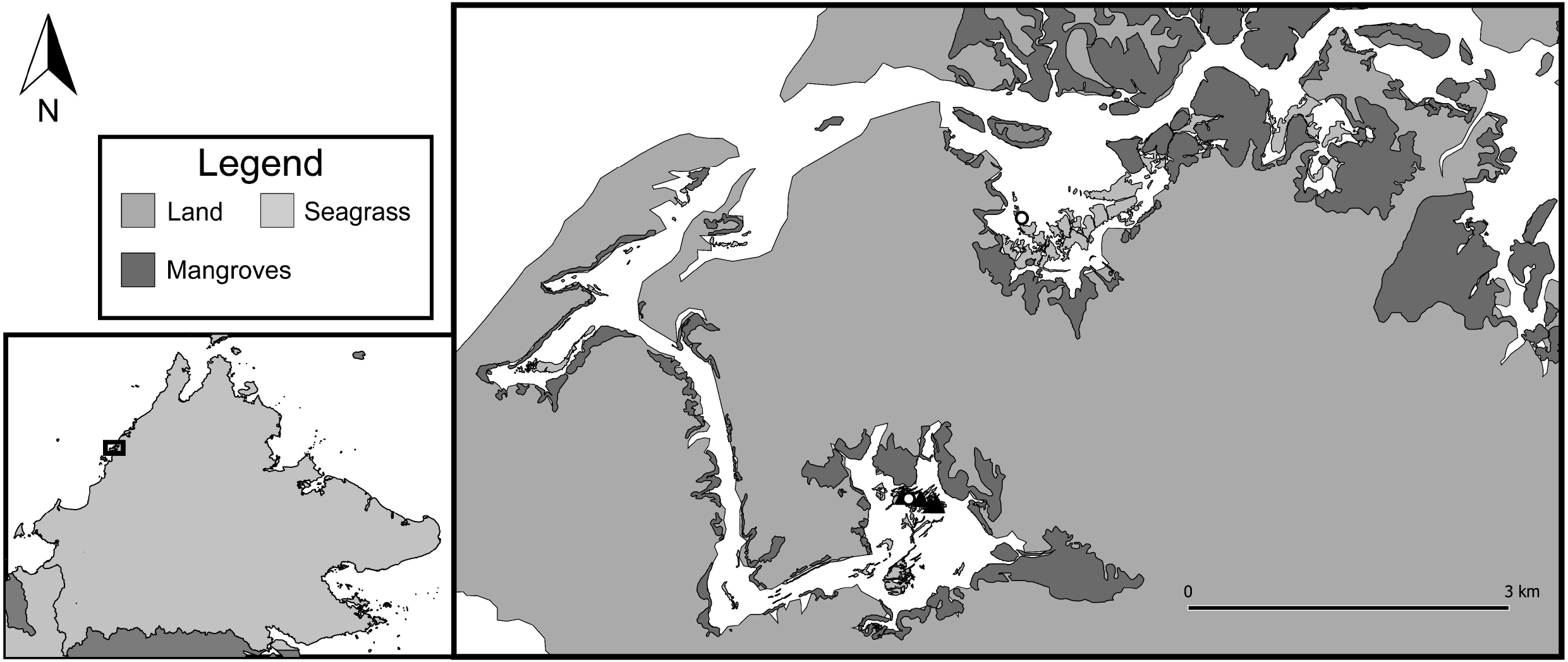
The Salut-Mengkabong estuary site used in the study. Salut is the southern arm of the estuary, while Mengkabong is the lagoon situated to the North. The sites at which the seagrass sediments were obtained for the incubation experiment are demarcated by black triangles (▲), while the sample cores used for SIT data are marked by white circles (○).The seagrass distribution information is based on collective indigenous knowledge, while the mangrove distribution is obtained from the World Atlas of Mangroves Version 3 (Spalding *et al*. 2018) and from Google Earth. (Map data: Google, 2019; Landsat/Copernicus, Digital Globe, Bornean Biodiversity & Ecosystems Conservation (BBEC) Sabah and WWF Malaysia, 2017.) The line map was produced with QGIS v3.6.0 and Adobe Illustrator CS6.

### Sediment collection and incubation

The sediments for the decomposition experiment were taken in 2016 from 22 cores from five sites in the estuary, spaced ~30 to 150 metres apart. The cores were transported back to the laboratory under ice (ambient temperature in icebox = 10.2 °C), where the surface 2 cm and 20–22 cm horizons were extracted and pooled. The latter horizon was taken a short distance ahead of the start of a transition to a lower, more fibrous brown facies (>26 cm). Samples from each sediment horizon were pooled in the manner of Westrich and Berner (1984) after wet sieving (1 mm) with previously filtered boiled seawater to remove large shells, debris and benthic fauna. After this, the pooled samples were divided into four separate Mason jars under nitrogen, and filtered boiled seawater was added to make up a 400 cm^3^ slurry with a final water content of 81.9%. Before the start of the incubation, the slurries were bubbled with N_2_ for 25 min and the anoxic status was checked (YSI ProDSS probe) before the Mason jar lids were replaced. To ensure that the sulphate supply was not limiting sulphate reduction, additional sulphate was added in stoichiometric proportion to the measured amount of CO_2_ emitted. This was done after the first month and again a further three times over the course of the first 300 days of the experiment. As a further precaution, sulphide and CO_2_ traps were placed in the jars’ headspace to both inhibit and control any build-up of metabolites and to measure net accumulative mineralisation. The sulphide traps were constructed by using epoxy to fasten a 110-mm-diameter Whatman No. 1 filter paper saturated with 1‰ zinc acetate to the underside of each jar lid. These were strategically folded to present a large total surface area and were placed alongside lead acetate paper strips to visibly detect any ongoing emissions of H_2_S. The filter papers were refreshed with fresh solution after every sampling procedure. The CO_2_ traps contained 2 to 3 g of dried high-absorbance-capacity soda lime (Dharmakeerthi *et al*. 2015) placed in 15 mL polypropylene centrifuge tubes. The tubes were open to the headspace and were replaced after each sampling time for further gravimetric measurements of CO_2_ accumulation rates. An additional set of soda lime traps was also placed in four Mason jars filled with filtered boiled seawater (400 cm^3^), which were added to the incubation cohort as CO_2_ procedural blanks (Keith and Wong 2006).

The Mason jar sediment slurries and blanks were all incubated at 30 °C in a constant-temperature room in the dark (covered in Al foil as a precaution against disturbance). The slurries were sampled after 7, 21, 42, 63, 105, 140, 175, 210, 308, 365, 400, 420, 470 and 500 days for POC, CDOM, ammonia, pH, and CO_2_. A Day 0 sample for POC was added after the first year. These were taken from the remaining pooled sediments (stored at −20 °C) and replicated with sediments from corresponding horizons within the sediment core used for the meadow’s geochronology. At selected times, samples were taken for δ^13^C_POC_, C:N_POC_ ratios for both horizons and ^13^CO_2_ trapped by the soda lime for the surface sediments.

After 500 days, additional aerated filtered seawater was added to the jars to bring the volume to back to 400 cm^3^ and the pH was adjusted to 8.5 with NaOH (Analar). The slurries were again kept in the dark at 30 °C and aerated for 30 days. To remove any possible organic and BC aerosols that might contaminate the slurry, the air was first passed through HEPA filters. The filters also supported a coarse polyester mat impregnated with charcoal. The pH of the slurry was adjusted every few days to maintain acidity between pH 7 and 8, and distilled water was added to replace any evaporative loss (Westrich and Berner 1984).

### Sampling and analysis protocols

The Mason jars were reopened under a N_2_ atmosphere and the pH of the slurry water was measured after the sediment had settled (ATC portable PH-107 (PH-009)), and their anoxic status was checked (YSI ProDSS). For sampling of the slurry, a cut-off syringe was used to extract 10 cm^3^ of slurry after thorough mixing; the subsamples were placed in 15 cm^3^ polypropylene centrifuge tubes and frozen at −20 °C before analysis. The remaining slurry was then bubbled with N_2_ for two minutes as a precaution to maintain the anoxic conditions within the jar. The lids of the jars were then resealed under N_2_ after the soda lime traps were removed, capped and replaced with identical traps. The traps were immediately oven dried and reweighed after first softly cleaning the surface of the centrifuge tubes of any accumulated red biofilm, and CO_2_ was determined gravimetrically (Keith and Wong 2006). Blanks indicated no significant leakage of air into the Mason jars and typically showed an increase in weight of ~0.0332 g (standard error (SE) = 0.014, *n* = 4), a value 68% less than the weight increase from the traps in the jars containing slurry samples.

After thawing, the slurry samples were centrifuged at 4000 rpm for 20 minutes to separate the pore water for measurements of CDOM_440nm_ (Harvey *et al*. 2015), ammonia (Strickland and Parsons 1968) and salinity (refractometer). The remaining sediment plug was then dried at 105 °C and the amount of water and sediment was noted to calculate the amount remaining in the mason jars for CO_2_ accumulation as dry weight of sediment after correcting for salinity (Lavelle *et al*. 1985). Particulate organic matter (POM), PIC and black organic matter (BOM) from the dried sediment slug was measured gravimetrically by loss on ignition (LOI_0.45g_) in a laboratory furnace (Carbolite CWF 1.8 L; Heiri *et al*. 2001; Chew and Gallagher 2018). Additional inter-batch corrections resulting from possible furnace aging and procedural handling differences were performed using in-house local sediment standards taken from the middle of the cores (*n* = 5) and randomly placed within the furnace. Standards were previously dried (60 °C) and stored frozen (−20 °C). All POM and BOM values were then converted to carbon content using a local calibration regression. The regression was constructed previously from sediments taken from Salut–Mengkabong seagrass and mangroves (Chew and Gallagher 2018) using the same furnace and in-house sediment standards. A coefficient of 0.273 used to transform the LOI_550-950°C_ to PIC by assuming the carbonate species to be calcium salt (Santisteban *et al*. 2004). However, it should be noted that a later analysis of the data suggested that the increase in carbonate may have been from ferrous salt. Until certainty is established, in both the form of thermal decomposition equation during the analysis and identity of the salt, all PIC contents are reported as CaCO_3_. All measurements are presented, except for CDOM_440nm_, in molar units for stoichiometric comparisons. CDOM_440nm_ was converted to DOC to give the organic dissolved pool dynamic an order of magnitude significance with other carbon variables. As far as we are aware, the calibration used for the conversion is the only one available for 440 nm determinations for an estuarine system (Harvey *et al*. 2015). The dataset is provided in the Supplementary Material should it be necessary for readers to rework the CDOM_440nm_ and PIC content in light of new information.

Analyses of stable POC isotopes of δ^13^C and their C/N ratios were performed on the two horizons across separate mason jars at selected times (Days 0 and 210). Before analysis, the samples were dried and vacuum sealed and sent to the Canadian Rivers Institute, University of New Brunswick Nature Laboratory (SINLAB). Re-drying after acidification (10% HCl (Analar)) to remove PIC was performed before analysis at the institute. No isotope or element analysis was done for the local source materials, which would typically be required for an estimation of their relative proportions. Nevertheless, estimations were gauged on the average ^13^C_POC_ and N:C endpoint signatures of seagrass, mangrove leaves and suspended particulate matter, using a model constructed for a number of tropical lagoons (Gonneea *et al*. 2004; Chen *et al*. 2017). In addition, stable isotopes of δ^13^CO_2_ trapped by the soda lime (days 7, 210, 308 and 500) were measured from a surface-horizon mason jar replicate. The jar was selected at random, and the analysis at the Central Science laboratory was performed by mixing ground samples and subsamples under Ar and placing ~2.5 mg into preflushed (Ar) vacutainers. The CO_2_ was released after dissolving the powder with pure phosphoric acid before injection. Handling errors were tested on one sample (mean, −19.78; SE, ±0.98, *n* = 4). Note that limited resources precluded any additional isotope analysis of either sediments or soda lime.

Sediment cores for the geochronology were collected using a sliding hammer Kajak corer (UWITEC, Austria) equipped with a 6 cm internal diameter polycarbonate core tube; the sediment–water interface was stabilised with a porous polyurethane foam plug. The core was transported vertically under ice to the laboratory for push extraction. Water content, bulk density, pore water salinity and loss on ignition at 550 °C and 950 °C were measured every 2 cm (Gallagher and Ross 2017). The remaining sediment for each horizon was used to determine particle size (laser diffraction, LSST-Portable, Sequoia, model: 220 Type B); after drying (50 °C) and storage for three months, ^210^Pb, ^226^Ra and ^137^Cs radionuclide analysis was performed using gamma spectroscopy at the Malaysian Institute of Nuclear Technology (MINT).

### Decomposition model

The reactivity continuum model was chosen to model the POC decomposition time series (Boudreau 1991; Arndt *et al*. 2013; Mostovaya *et al*. 2017). Exploratory analysis indicated that this gave the best fit and was the most parsimonious descriptor of the POC dynamics over single and multi G models (Arndt *et al*. 2013). The model fits a continuous distribution of organic matter decomposition, from labile to increasingly recalcitrant, and was calculated as follows:

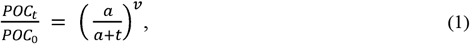

where *a* is the apparent age of the organic mixture (years) within the deposit, as a measure of its degradability relative to an apparent age at the time of deposition. The exponent *v* is the gamma distribution coefficient, which describes the labile–recalcitrant distribution and dominance (1 to 0, respectively) of the sediment horizon’s organic mix. Taken together, the initial first-order decomposition coefficient, *k*_0_, is defined as *v/a*, which becomes increasingly recalcitrant with incubation and burial time *t*. The parameter solutions were calculated iteratively using a nonlinear least squares parameter estimation within the platform SigmaPlot 12.0. It should be noted that there is a rival continuous diagenetic model. The model, ostensibly constructed within phytoplanktonic- and bacteria-dominated sediments, uses a power function to describe how organic matter becomes increasingly recalcitrant over apparent time (Middelburg 1989). While the two models are equivalent mathematically (Tarutis 1993) when applied within closed systems such as jars (i.e., no sediment accretion), the sediment’s mix of seagrass litter, microalgae and mangroves (see Results), with very different intrinsic reactivities (Middelburg 1989; Kristensen 1994), would seem more aligned with an RC explanation than a relatively less parsimonious power model as a sum of differing degrees of aging across different organic sources.

### Geochronology

Sediment isotope tomography (SIT) was used to model a continuous ^210^Pb geochronology down the sediment core’s uninterrupted depositional regions (Gallagher and Ross 2017). The model describes how the ^210^Pb activity of sedimentary horizons can be fitted to a function that includes the changes in the ^210^Pb flux and sedimentation velocity as the ^210^Pb decays over time (Carroll *et al*. 1999). The algorithm employs a parsimonious inverse solution to best simulate the ^210^Pb profile by solving for the model’s parameters for maximum disentanglement of the flux and sedimentation velocity terms (Liu *et al*. 1991). Further constraints and evaluations of solutions can be made by the presence of known events (Carroll *et al*. 1999). Such events are traditionally peaks or horizons of ^137^Cs from atomic fallout within baseline sediments and depositional facies characteristic of surrounding material brought in by storms, earthquakes, floods or tsunamis.

Supporting data, additional figures and method details can be found in the electronic Supplementary Material.

## Results

### Sediment core descriptions

The first 23 cm of the Salut and 25 cm of the Mengkabong meadows were visibly muddy (black) with no evidence of bioturbation. Below the 23 and 25 cm horizons, the character of the sediment visibly changed to a coarser mixture of more compact light and dark brown sediments containing a plethora of shell and mangrove wood debris (refer to supplementary Fig. S2). No sulphide could be detected by smell or with lead acetate strips left in the sediment for a minute while they were extruded into receiving tubes before separation.

### Sediment horizon organic composition

The ^13^C_POC_ and their N:C ratios taken through the incubation did not appear to change and the two horizons exhibited near identical signatures (Table 1). These signatures converged even further when the effects of diagenetic transformations were considered (Galman *et al*. 2008; Galman, Rydberg *et al*. 2009). Interestingly, it was found that seagrass litter was likely a minor component (around 5%). The remaining components of surface-suspended matter, ostensibly microalgae, and mangrove sources made up the remaining 25% and 70% respectively (refer to Supplemental Material), in agreement with other ecosystems in the region (Chen *et al*. 2017).

**Table 1:** Dry mass of particulate sedimentary carbon and stable nitrogen isotopes and their molar ratios from the incubation jars. S and B refer to the surface (0–2 cm) and bottom (20–22 cm) horizons, followed by the day number during the incubation on which the sediments were extracted. All δ^13^C values have been normalised to preindustrial times (Suess effect) using their modelled depositional age. S0 and B0 are from single samples, while S210 and B210 are the means of four subsamples with their respective standard errors.

| Sample | $\delta^{13}\text{C}$ (‰) | C (%) | N (%) | N/C ratio |
| --- | --- | --- | --- | --- |
| S0 | –24.61 | 7.83 | 0.61 | 0.066 |
| S210 | –24.71 ± 0.04 | 7.64 ± 0.13 | 0.58 ± 0.004 | 0.065 ± 0.0007 |
| B0 | –24.06 | 7.47 | 0.62 | 0.071 |
| B210 | –24.22 ± 0.03 | 7.61 ± 0.11 | 0.63 ± 0.004 | 0.070 ± 0.0007 |

### Geochronology

While the depth of the storm facies were similar, it was clear from the ^210^Pb activity profiles that the sedimentation dynamics within the baseline sediments were very different. The Salut meadow, an embayment isolated at the head of the branch and fed by a rivulet, supported peaks in activity at around 10 cm (Fig. 2), in contrast to a general decay in ^210^Pb activity from the surface of the Mengkabong meadow (Fig. 2), an embayment isolated from the main branch. The difference in dynamics was also highlighted in the inability to detect any ^137^Cs activity from atomic fallout events within Salut sediments, which were evident as significant ^137^Cs activity between 5 cm and 13 cm, peaking at 5 cm down the Mengkabong meadow core. This relatively shallow signal is consistent with blow back of fallout from the 2011 Fukushima Daiichi nuclear accident (Kaeriyama 2017).

**Fig. 2.**
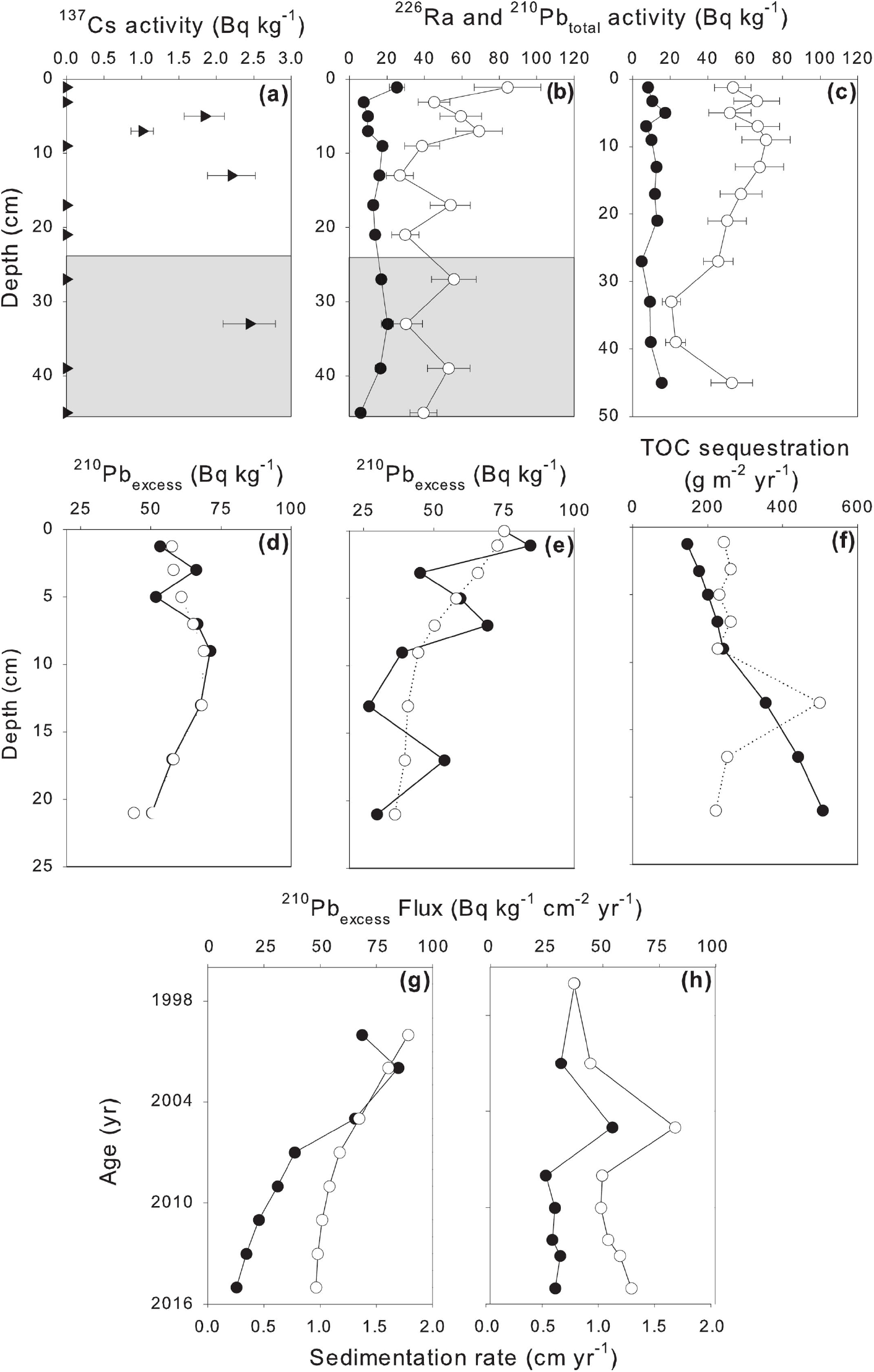
Radiogeochronological profiles down the upper seagrass sediments of Salut-Mengkabong estuary/lagoon. The shaded area represents the mangrove deposition event. (*a*) The ^137^Cs activity (►) down the Mengkabong meadow sediments; no activity could be detected down the Salut meadow sediments. (*b, c*) The respective supporting ^226^Ra (●) and total ^210^Pb_total_ activity (○). (*d, e*) The resultant mean excess or unsupported ^210^Pb_excess_ activity, corrected for ^226^Ra, outside the deposition event (●) superimposed on their respective stable sediment isotope tomography (SIT) simulations (○), together with (*f*) their resultant POC sequestration rates for Mengkabong (○) and Salut meadows (●). (*g, h*) Changes over time in the sedimentation and ^210^Pb_excess_ parameters as simulated by SIT in the Mengkabong and Salut sediment columns, with black circles indicating the actual recorded ^210^Pb_excess_ activity and white circles indicating the ^210^Pb_excess_ activity as modelled by SIT.

When the SIT solution for the Mengkabong system was constrained by the timing of the Fukushima fallout, the depositional event’s age was estimated as ca. mid-1990s. The only recent weather event of note was from the passage of Tropical Storm Greg (December 1996). The storm is regarded as a once in 400 years occurrence for this region, which is commonly known as ‘The Land below the Wind’ due to its location south of the influence of the Typhoon Belt. The 1996 storm triggered floods that severely affected the west coast of the state (Abdullah and Tussin 2014), and a local resident shared his experience as a witness to a coastal surge of ~4 m within the adjacent mangrove forests (Mohd. Asri Mohd. Suari, personal communication). With the confirmation that the depositional event was likely to be Tropical Storm Greg, the SIT model now adds constraints for the Salut meadow baseline sediments of age no older than 1996. Based on these solutions, the origin of the very different ^210^Pb dynamics becomes apparent. In Salut, both the flux of the meadow’s excess ^210^Pb activity and the sedimentation rates fell over time. In Mengkabong, rates of sedimentation and ^210^Pb activities remained relatively constant (220 g m^−2^ per year, Fig. 2) and were only interrupted by an increase consistent with shoreline development during a peak in annual rainfall (ca. 2005, unpublished data). These show relatively high sequestration rates near the top of the range, even before any correction for loss over time (Fig. 2).

### Incubation experiment

Throughout the incubation experiment, the pH of both surface sediments and sediments taken from 20–22 cm became increasingly acidic over time (Fig. 3). The older sediments taken from 20–22 cm were more acidic and remained invariant and acidic. Surface sediment slurries, in contrast, were initially less acidic; however, their acidity increased over time, reaching an asymptote after 300 days equal to that of the older sediment slurry. The experiment failed to detect the presence hydrogen sulphide within the jar’s headspace (no blackening of the lead acetate strips) that would infer ongoing sulphate reduction.

**Fig. 3.**
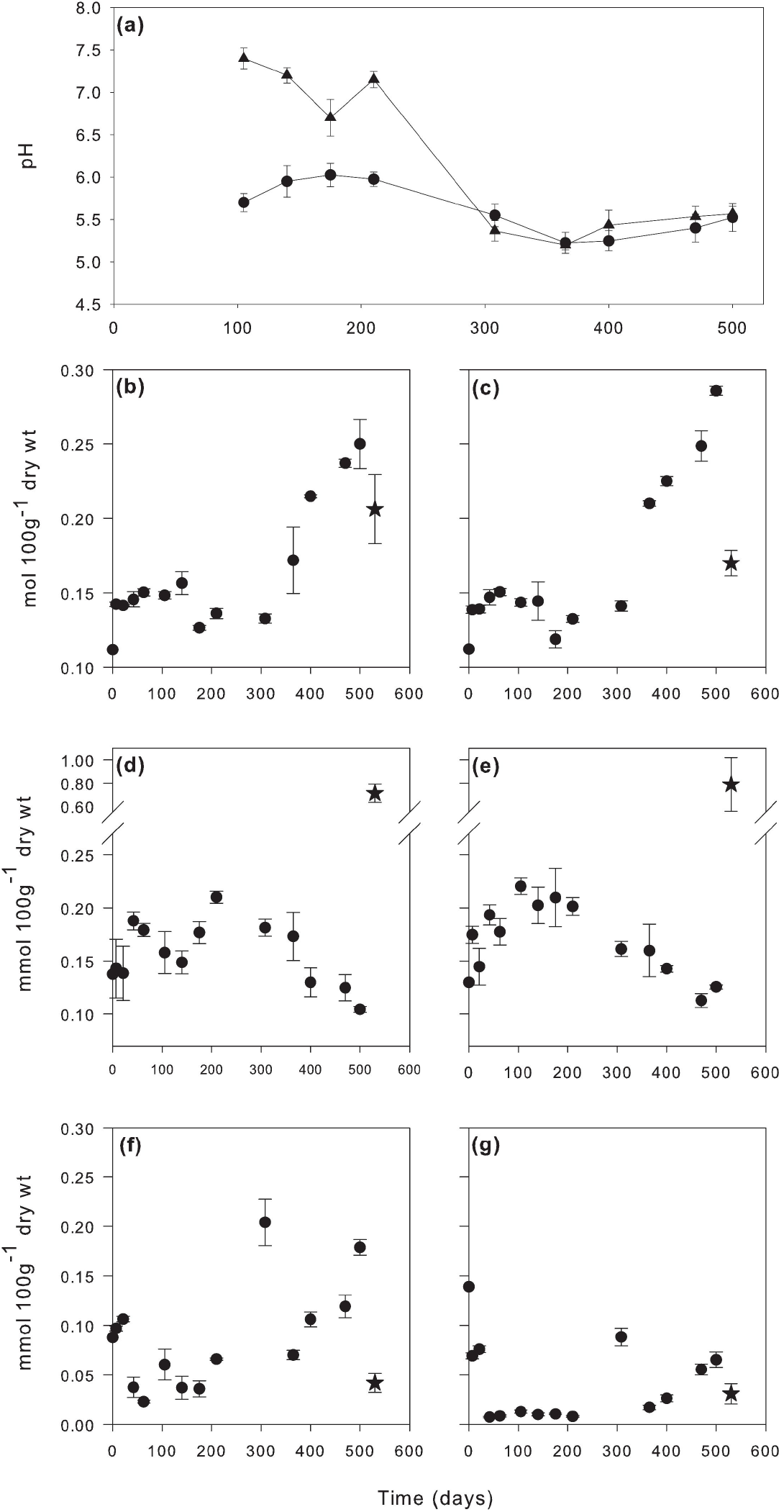
Values of pH, PIC, DOC and ammonia measured in the sediments throughout the incubation experiment. (*a*) The pH of the sediment slurries from day 105 until the end of the anoxic incubation period. (*b, c*) The PIC content of the sediment. (*d, e*) The DOC content of the porewater of the sediment slurry. (*f, g*) The ammonia concentrations of the porewater of the sediment slurry. (*b*), (*d*) and (*f*) correspond to the surface 2 cm horizon, while (*c*), (*e*) and (*g*) correspond to the sediment collected from the 20–22 cm horizon. The last point in each series, indicated by a star (⃬), shows the final values of the sediments after a 30-day reoxygenation period. Error bars indicate standard error (*n* = 4).

While the initial BC represented a modest fraction of the POC (0.079 and 0.067 mole/100 g or 11% to 13%), its influence on the POC dynamics was not apparent as there was no significant decay in the BC content over the 500 days, and RC solutions with the time series failed to converge. The anoxic decay of POC for the surface and older 20–22 cm horizon sediments fitted the RC model well, and the separation of the terms was within acceptable limits (Fig. 4). Surface sediment POC content was greater than sediments taken from 20–22 cm. However, we found no significant difference in their RC decay and apparent age parameters for the decomposable fraction (Fig. 5) despite different interdecadal depositional ages (18 years). Projections suggested that both horizons would have lost close to 30% of their POC content within the first several years (6 to 7)^1^. Nevertheless, the overarching dynamics were such that both horizons converged to losses of around 49% and 51% after 100 years of burial.

**Fig. 4.**
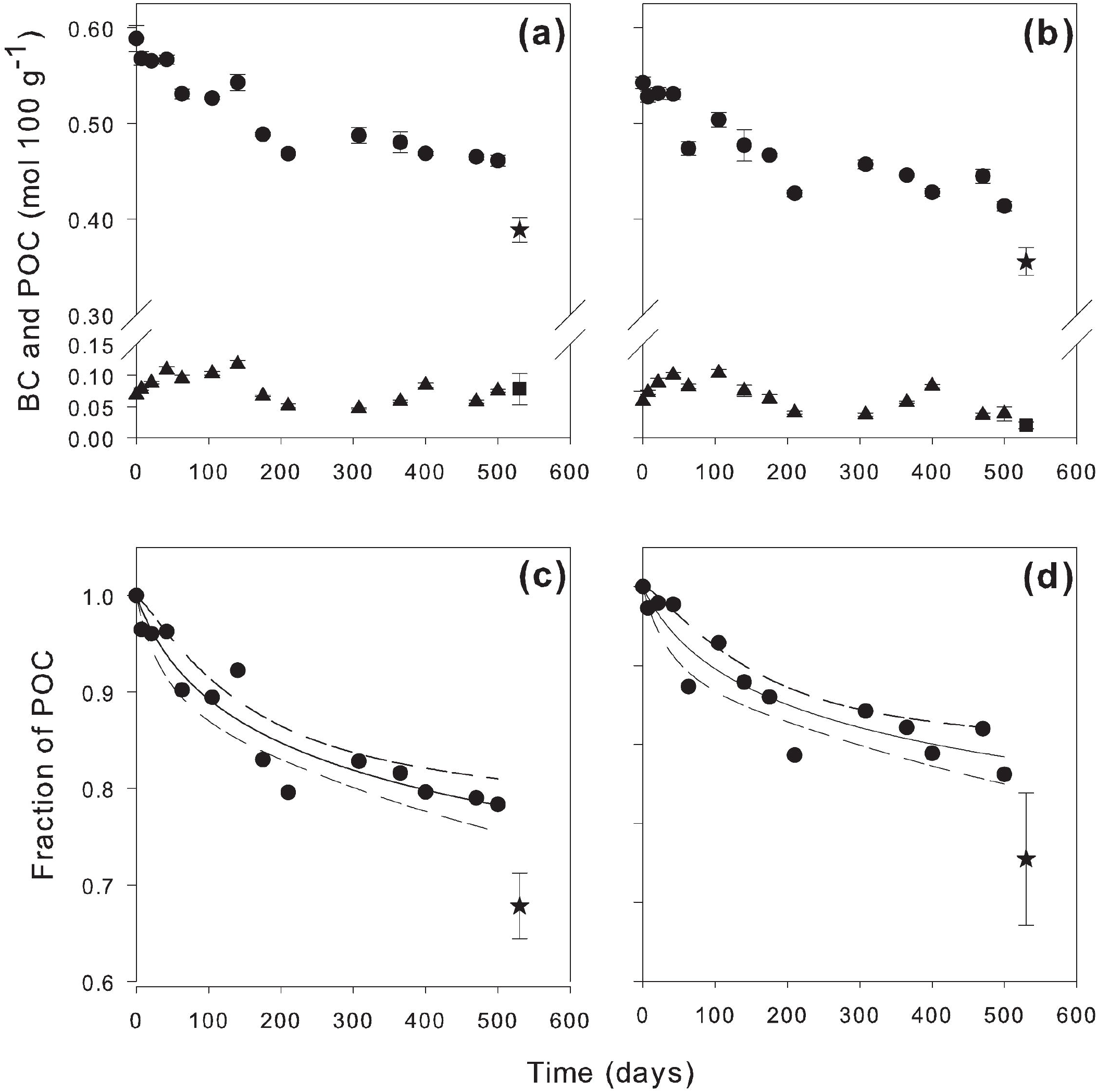
Particulate organic matter (POC) content, Black Carbon (BC) content and loss of POC fraction of the sediments used over the anoxic incubation and subsequent reoxygenation. The mean POC content, corresponding to (*a*) the surface 2 cm and (*b*) the sediment collected from the 20–22 cm horizon, is shown by the series marked by circles (●). The mean BC content is shown by the series marked by triangles (▲). Error bars indicate the standard error (*n* = 4). The loss of the POC fraction over time in (*c*) the surface 2 cm and (*d*) the sediment collected from the 20–22 cm horizon using the reactivity continuum model. Broken lines indicate the 95% confidence limit, as do the errors on the final point. The last point in each series, indicated by a star (⃬) for the POC series and a square (■) for the BC series, shows the final value of the sediments after a 30-day reoxygenation period.

**Fig. 5.**
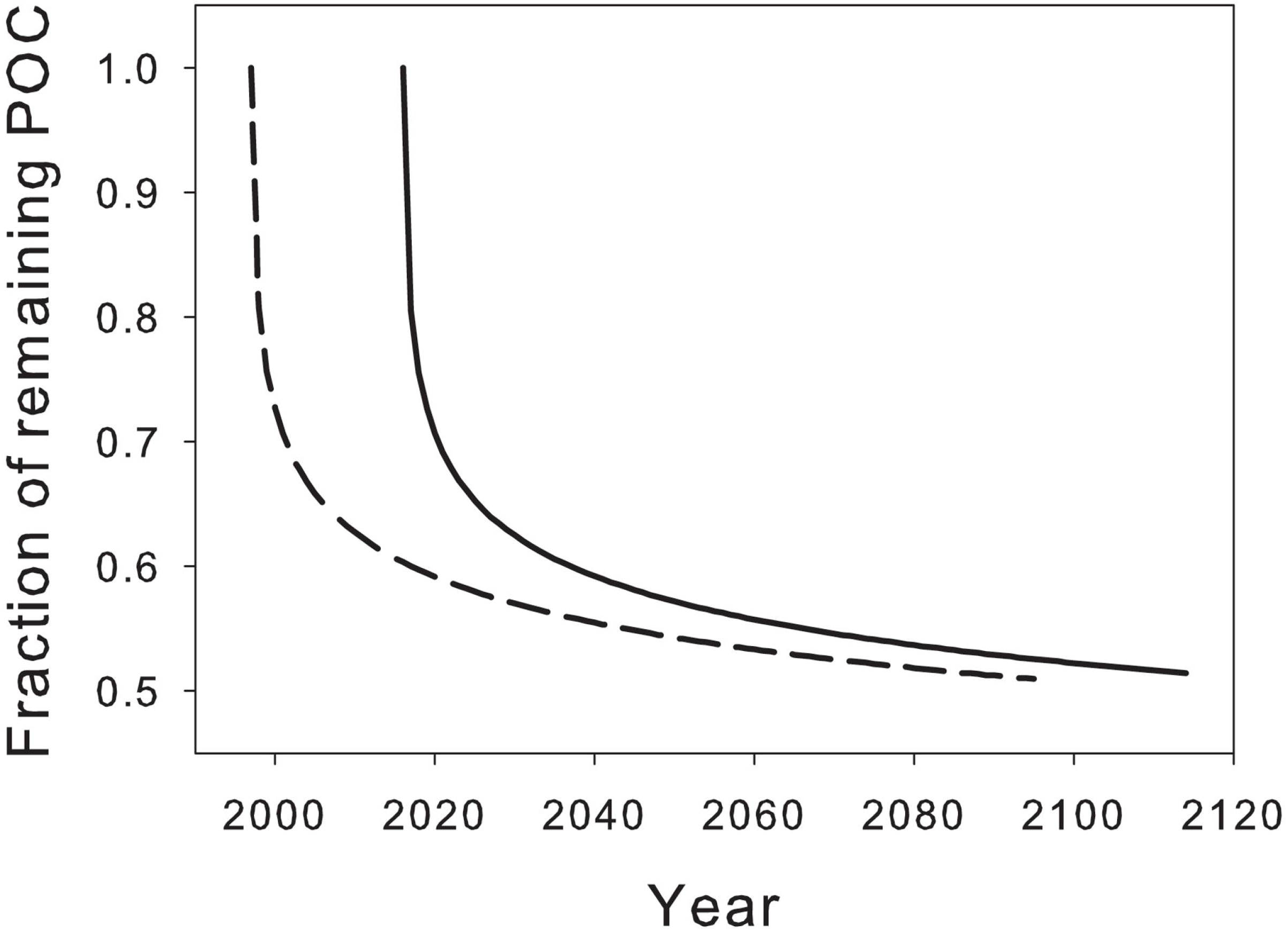
Extrapolations of the fraction of remaining POC within the sediments over 100 years following deposition. The broken line corresponds to the sediments collected from the 20–22 cm horizon, which were dated to deposition circa 1996, while the solid line corresponds to the sediments collected from the surface 2 cm, deposited in 2016.

In contrast to POC, the dynamics of PIC, DOC and NH_3_ were not continuous. After around 300 days, the carbonate content started to increase for both sediment horizons and appeared to move toward an asymptote. This was accompanied by an increase in ammonia and a decrease in DOC content (Fig. 3) after the ammonia content had first fallen and the DOC content increased (Fig. 3). DOC and ammonia pools were notably an order of magnitude smaller than POC. Only the cumulative CO_2_, after correction for PIC generation after the 300 days, showed steady-state dynamics which slowed towards an asymptote (Fig. 6). However, there appeared to be a notable deficit in the amount of CO_2_ emitted for the amount of POC decomposed, in particular for the deeper, older sediment horizon. Furthermore, the δ^13^C_POC_ isotopic signatures were not coupled to each other. The δ^13^CO_2_ values extracted from the soda lime were both relatively constant and very much heavier and relatively constant than the POC source mix. This was −19.78 ± 1.95 (*n* = 4) at Day 7, −17.74 (*n* = 1) at Day 189, −19.30 (*n* = 1) at Day 308 and −18.56 (*n* = 1) at Day 500, the end of the incubation experiment. Meanwhile, at the same time, the ammonia, DOC and PIC contents in the sediment slurry remained relatively constant up until around day 365, when a change in trend was observed (Fig. 3). From Day 365 until the end of the incubation experiment, both PIC and ammonia levels in the surface sediment slurry increased, with an increase of 46.48% (SE = 3.91, *n* = 4) in PIC and 60.86% (SE = 1.57, *n* = 4) in ammonia levels, while DOC levels dropped by as much as 73.77% (SE = 8.75, *n* = 4) over the same period of time. Meanwhile, for the sediment slurry taken from 20–22 cm, PIC and ammonia levels increased by 50.57% (SE = 1.44, *n* = 4) and 73.19% (SE = 2.17, *n* = 4), respectively, while DOC levels dropped by 28.44% (SE = 4.89, *n* = 4).

**Fig. 6.**
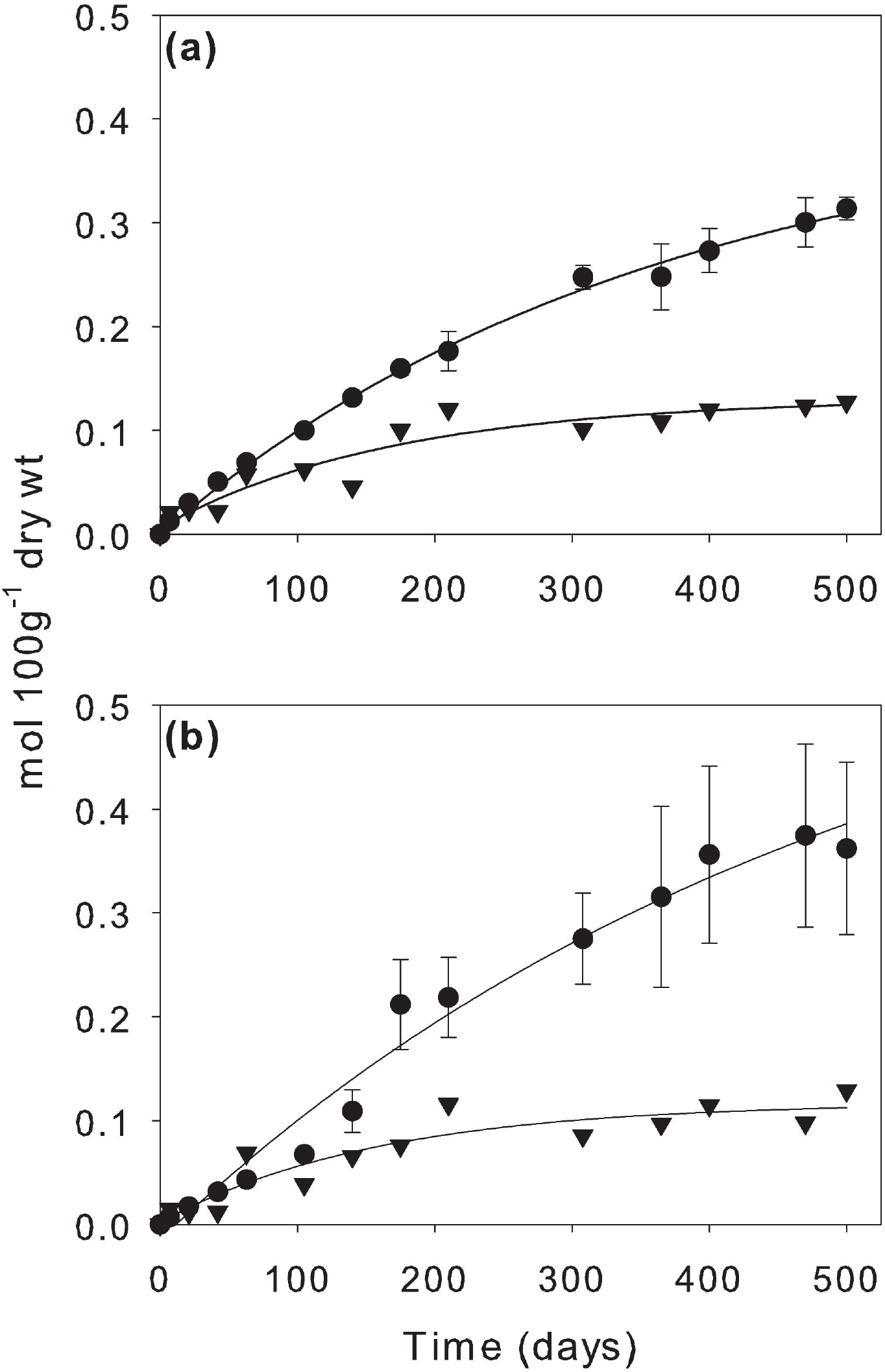
Cumulative CO_2_ absorbed by soda lime and net loss of the POC fraction of the sediments used over the anoxic incubation. (*a*) and (*b*) correspond to the surface 2 cm and the sediment collected from the 20–22 cm horizon, respectively. Error bars indicate the standard error (*n* = 4). The series indicated by circles (●) is the cumulative CO_2_ absorbed over the course of the incubation, while the series indicated by triangles (▲) is the cumulative loss of POC over the same period.

### Aeration incubation

The short aeration pulse over 30 days after the completion of the 500-day anoxic incubation showed a large decrease in POC (18.86%, SE = 4.09, *n* = 4 for surface sediments; 16.99%, SE = 5.04, *n* = 4 for sediments from 20–22 cm), outside that of the parameters of the anoxic mineralisation models (Fig. 4). This increase in decomposition was also in line with a disproportionate increase in DOC over the anoxic mineralisation, confirming that for both horizons, organic aging had little effect on the recalcitrance of the buried POC.

## Discussion

### Decomposition

Assuming the incubation was sufficiently long to capture interdecadal decay parameters, it appears that POC deposited, on average within one to two years of deposition may suffer significant losses over climatic scales (49% to 51%). However, we must suggest caution in applying the surface horizon extrapolations as a generalisation to seagrass beds in other locales, as such sediments will inevitably change their redox status from an aerobic to an anaerobic dominated form of mineralisation. Aerobic mineralisation is clearly the more rapid of the two, the result of greater efficiency in the mineralisation of the recalcitrant fractions (Kristensen *et al*. 1995). As well as changing redox conditions, the nature of the organic mixture will likely affect the decay parameters of the RC model. Clearly, the remaining half of the organic carbon, a seemingly recalcitrant fraction, is more than can be accounted for by the <10% contribution of the BC alone. It is also unlikely in this case that any presence of phytolith occluded carbon was responsible given that the BC methodology may have inadvertently included this form (Chew and Gallagher 2018). What remains is up to speculation; it may consist of bacterial necromass (Burdige 2007) and—an increasingly important vector, especially within Southeast Asian coastal ecosystems— microplastics (Nor and Obbard 2014; Li *et al*. 2019). While microplastics have turnover times of over 1000 years (Gewert *et al*. 2015), their amounts as carbon within soils and sediments remain largely unknown. Some values have been estimated for terrestrial soils (Rillig 2018), ranging from 0.1–5% of POC for pristine environs to as much as 6.7% by soil weight.

Whatever value the overall decay parameters may take over space or time, it remains puzzling that we found little difference in the POC decomposition model parameters between the surface and the deeper sediment horizons. This was not apparent in coastal non-vegetative sediments, which are dominated by more labile phytoplanktonic organic sources (Burdige 1991; Zimmerman and Canuel 2002). This can be explained by two possible theories: either the sediments in these types of meadows were well mixed, which is unlikely given the presence of ^137^Cs peaks and ^210^Pb decay series, or the stable isotope signatures and recalcitrance are not covariant down the sediment columns. For the latter to be consistent, mangrove sources would need to balance an increase in recalcitrance between or within other organic sources as they are buried over time. In essence, a mix of the reactivity continuum and power models would best describe this. However, it cannot be discounted that changes in physical protection and benthic consumption parameters may also play some role (Arndt *et al*. 2013).

### Diagenesis and the coupling between CO_2_ and decomposition

The mineralisation and decomposition series have several notable features. These are seemingly punctuated dynamics of carbonate, ammonia and DOC, the CO_2_ deficits with POC decomposition, and the notably heavier ^13^CO_2_ signature over that of ^13^C_POC_. These dynamics suggest that the incubation experiment was not at a steady state as different diagenetic processes switched on and off. How this affects the decompositional model’s parameters is uncertain, but it is unlikely that the result is an underestimate, given that the observed diagenetic switches likely reflect a resource limitation that the incubation failed to supply. Nevertheless, this limitation is common to any natural perturbation experiment attempting to discover what is possible under a different set of conditions than that which may be encountered in other systems.

Within the limits of our monitored variables, the results imply that the initial fall in ammonia content under dark anoxic conditions is synonymous with coupled dissimilatory nitrate reduction (DNRA) and denitrification by anammox autotrophic CO_2_ fixation (Ni and Zhang 2013). Indeed, recent work has also shown an unexpectedly high degree of anammox and DNRA in the upper muddy seagrass sediments of a subtropical lagoon (Salk *et al*. 2017). Nevertheless, the relatively small changes in NH_3_ indicate that any dark CO_2_ fixation would not have affected the overall CO_2_ dynamics, even after considering a stoichiometry of C:NH_3_ of 15:1 (Koeve and Kähler 2010). Although, it could be argued that the production of archaeal necromass may have contributed to an increasingly recalcitrant pool of POC over time (Burdige 2007), the extent to which this would contribute to the decomposition dynamics would depend, in part, on the supply of nitrate for coupled DNRA. A reduction in the supply of nitrates may perhaps be responsible for a change to another mineralisation process responsible for the increase in both NH_3_ and PIC after 300 days.

Anoxic PIC and NH_3_ production within marine coastal sediment, while consistent with sulphate reduction (Burdige 1991; Mucci *et al*. 2000), is also inconsistent with several sedimentary parameters and observations. First, we could not detect any H_2_S produced within the Mason jar headspace throughout the incubation period. Second, molar NH_3_:CO_2_ ratios were clearly an order of 10^3^ larger than those found for marine sediments dominated by sulphate reduction (Burdige 1991). What is not clear are the reasons for the increase in PIC, of sufficient amounts to affect the CO_2_ dynamics. Nevertheless, the lack of evidence for significant levels of sulphate reduction and alkalinisation points to another type of mineralisation, one that can support a suitable acidic microenvironment. Recently, it has been demonstrated that an iron-reducing bacterium can precipitate siderite (FeCO_3_) within acidic sediments at ambient temperatures (30 °C). It was suggested that alkalinisation at the cell walls was induced mainly by its production of NH_3_. Indeed, the dynamics of the parameters measured herein fall within the scientific justification of inference to the best explanation (Lipton 2000). The sediments were acidic and there was a parallel rise in NH_3_ production with PIC outside the stoichiometry of sulphate reduction. Furthermore, additional analysis of selected remaining sediment samples retained throughout the incubation experiment indicated that the total iron content was sufficient to support siderite formation (0.051 mol 100 g^−1^, SD = 0.0064, *n* = 60; see Supplementary Material), but only to levels to which the carbonate appears to be reaching an asymptote (~0.15 mol 100 g^−1^, Fig. 3).

What is clear, however, is that the overall CO_2_ dynamics observed fall well short of accounting for the continued loss of POC irrespective of PIC and DOC. By itself, this implies that there must be another mineralisation product. As far as we are aware, methane formed from methanogenesis is the remaining alternative. Methanogenesis would result in the release of both CO_2_ and CH_4_, within its own sedimentary niche, where any iron reducers cannot directly compete (Bray *et al*. 2017). While we did not measure methane during this incubation, the supposition is supported by relatively constant ^13^CPOC values and considerably heavier ^13^CO_2_ ratios over the incubation (Table 1 and 2). Such patterns have also been found for highly organic coastal marine sediments where a considerably lighter ^13^CH_4_ (~58.9‰) balances out the heavier ^13^CO_2_ (~19.2‰) fraction, to maintain a constant heavy source of ^13^C_POC_ over time (Boehme *et al*. 1996). Why methanogens should dominate mineralisation over sulphate reduction is not clear. Perhaps it is due to the high acidity of sediments seemningly supplied from the adjacent mangrove mudflats (Marchand *et al*. 2004) and iron reducing bacteria (Koschorreck 2008).

## Conclusions

The incubation experiment appears to capture the long-term decomposition parameters for POC. The RC model seems to indicate that current estimates of carbon sequestration may be significantly overestimated, in this case by around 50%, unless corrections can be made for loss over centennial time scales. Furthermore, much remains to be investigated on the coupling of POC losses to greenhouse gas emissions that have different atmospheric warming effects and the roles of processes post disposition, such as dark CO_2_ fixation and carbonate formation on net CO_2_ emissions. Without certainty in both the estimates and the conceptual model, there will not be sufficient certainty in the estimates of carbon storage and sequestration services rendered by seagrass ecosystems for use in cap-and-trade carbon markets to embrace these ecosystems as part of a solution to climate change.

## Author contributions

C.C.H. and J.B.G. assisted in fieldwork and design of equipment and analysis of the iron content. C.C.H. carried out the incubation experiment and the remaining analysis variables, created the figures and tables, compiled the supplemental material and the statistical analyses within the tables and contributed to the modelling. J.B.G. was responsible for the concept, the final modelling solution and led the writing of the manuscript. C.S.T. collected cores and performed the SIT ^210^Pb event geochronology under supervision from J.B.G. N.M.Z. provided the statement on recalcitrant carbon in the form of micro plastics found in the discussion. All authors approved the final version of the manuscript and agree to be accountable for all aspects of the manuscript.

## Acknowledgments

Our thanks go to our boatman Awang Azmee and Michael Yap Tzuen-Kiat for help in collection and analysis of the samples.

## Footnotes

1 The 30% was calculated as the time of symmetry of the decay series second derivative as percentage lost over percentage of time over a span of 100 years (Δ*lost*/Δ*t* = 1). Although it is a continuous function, as both scales are of the same magnitude, it thus marks the threshold time of a significant slowdown in decomposition.

