## Supplemental Material for "Blue carbon sequestration dynamics within tropical seagrass sediments: Long-term incubations for changes over climatic scales"

**Collection and processing of sediments used in incubation experiment**

The sediments were taken from 22 cores from 5 different sites within the northern part of the Salut estuarine lagoon, spaced around 30-150 metres apart. Samples were collected with an acrylic corer with core lengths from 37 cm – 47 cm cores. The sediment cores were taken by the boatman, Mr. Awang Azmee, by hand, and capped with rubber bungs. The core was placed in a bucket vertically and the sediment that was resuspended by the coring action in the water column of the corer was allowed to be deposited back onto the surface of the sediments before extrusion. An extruder bar with 0.5 cm graduations was used to push in one of the smaller bungs from the bottom into the tube, thus pushing out the sediment column from the corer tube. The top 2 cm of the sediment column was first cut off from the sediment column with a plastic cake knife then immediately bagged in sealable Ziploc bags and placed into an icebox (ambient temperature in icebox = 10.2 °C). This process was also repeated for the 20-22 cm section of the cores, being stored in a separate Ziploc bag. The rest of the sediment column was discarded, with the exception of the 2-20 cm section of 5 cores, which were also kept and taken as an in-house standard to ensure accuracy during the loss on ignition phase of the experiment between different batches. Sediments from the same sections of different cores were placed into the same Ziploc bags until they were half full, after which the bags were replaced. The Ziploc bags were squeezed around and mixed in order to homogenize the sediments from the different cores. Once the sediments arrived back at the laboratory, the bags were placed into a fridge at 7 °C and stored overnight. The sediments were then sieved while wet using a sediment shaker with a mesh size of 1mm, in order to remove any large particles and organisms which may cause biases due to more organisms living at certain depths. Any benthic organisms growing in situ would also not constitute the total organic matter protected from remineralisation as well, and thus their removal is justified. The sediments were mixed into a slurry and placed into the glass mason jars, ensuring that the same amount of water content was present in each of the jars. The total volume of the slurry in each jar was 400 ml, with the sediment from 20-22cm having a 1:1 ratio of sediment to seawater, and the sediment from 0-2 cm having its sediment:seawater ratio adjusted accordingly to match, giving both slurries a water content of 81.9%. The seawater used was obtained during high tide at the UMS Jetty on the 19th of October 2016 at 15.37, filtered through Whatman GF/B filter paper and boiled to remove any microorganisms and dissolved gases.

**Table S1:** Coordinates of the sites where the cores used in the study were taken. Sites starting with ICB denote the coordinates of the 5 sites where the cores for the incubation experiment were taken from.

| **Site** | **Longitude (° N)** | **Latitude (° E)** |
|---|---|---|
| Salut SIT | 6.10728 | 116.15278 |
| Mengkabong SIT | 6.13083 | 116.16228 |
| ICB 1 | 6.10745 | 116.15231 |
| ICB 2 | 6.10727 | 116.15382 |
| ICB 3 | 6.10670 | 116.15473 |
| ICB 4 | 6.10667 | 116.15525 |
| ICB 5 | 6.10688 | 116.15517 |

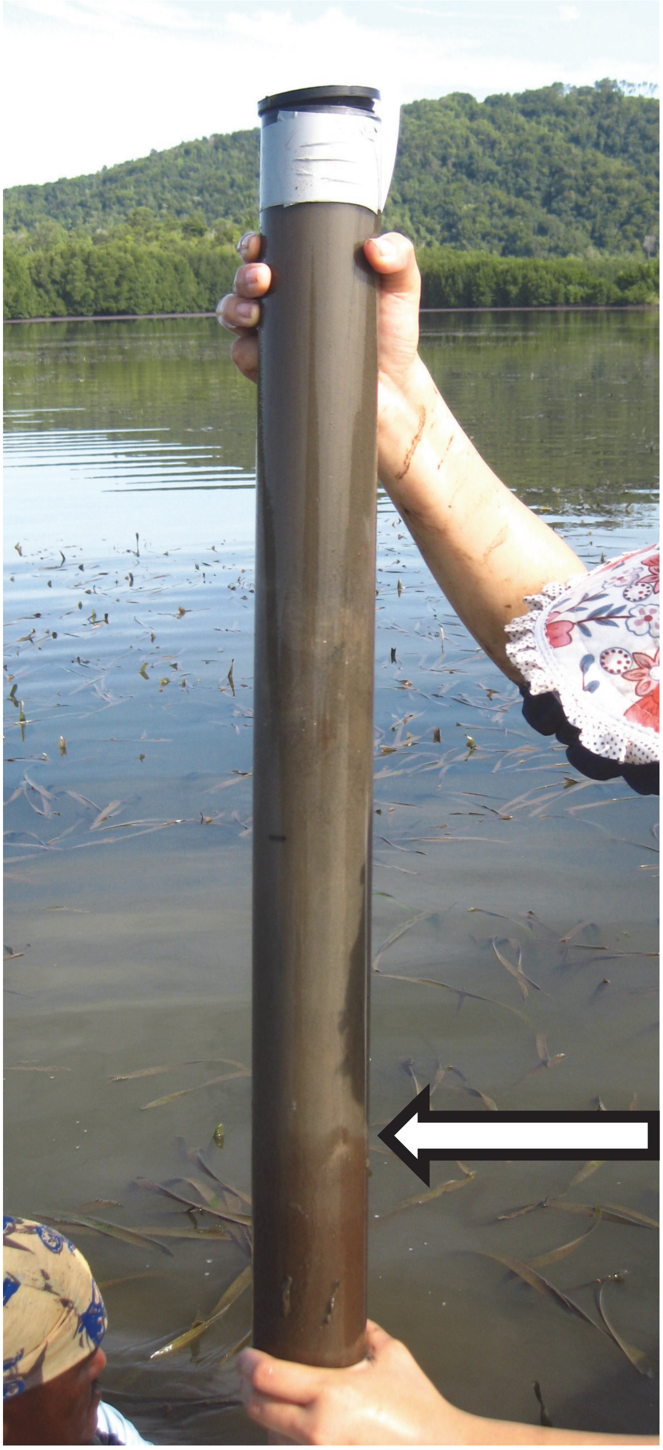

**Supplemental Figure S1:** Sample of one of the sediment cores taken for the incubation study at Salut Lagoon. Note the transition in the colour of the sediment at ~24cm, indicated by the arrow.

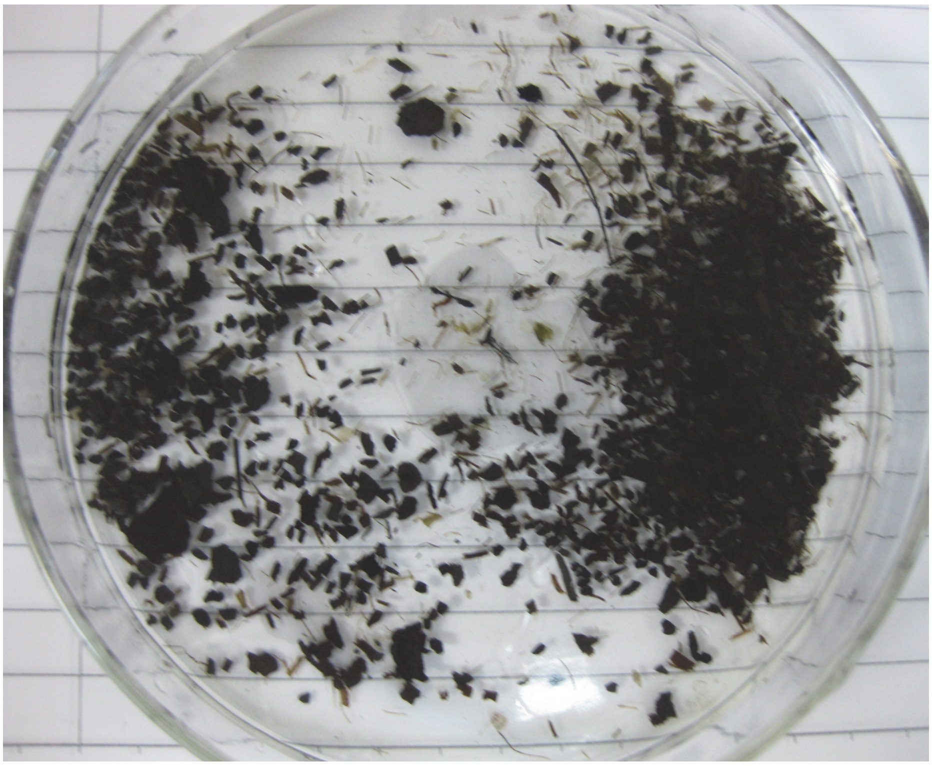

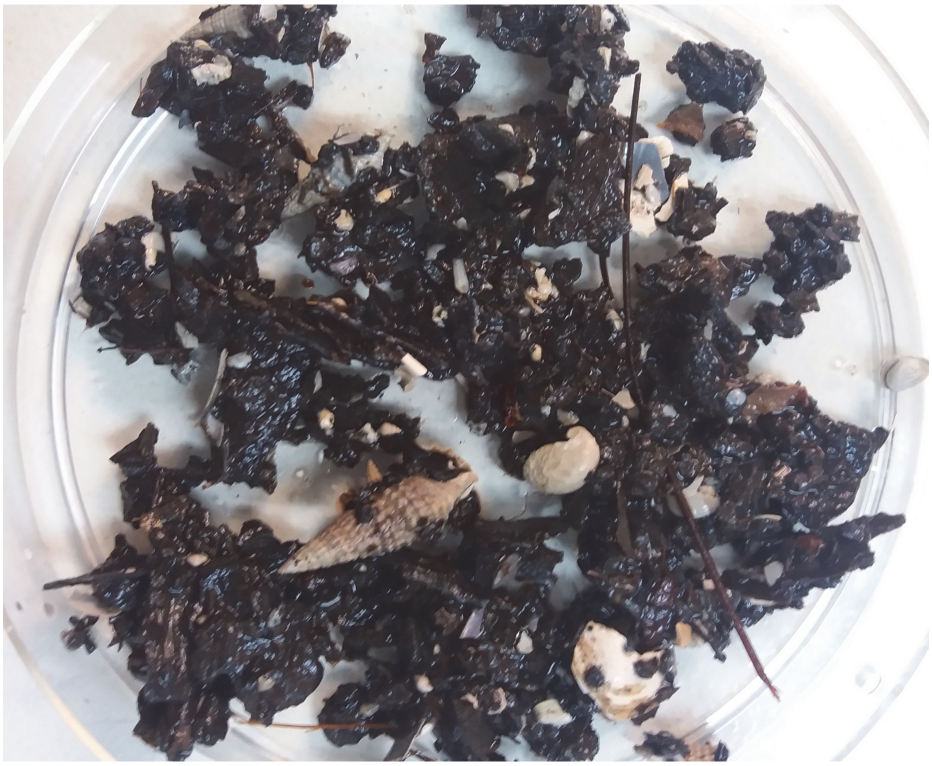

**Supplemental Figure S2:** The remnants of detritus larger than 1mm in size retained on the sieve after wet sieving. Top: Surface 2 cm sediment detritus, Bottom: Detritus taken from below 26 cm depth, within the lower, more fibrous brown facies. Both sediments were taken from the seagrass beds within the northern Salut estuary.

**Incubation experiment design and methods**

The experiment makes use of 1 litre glass mason jars, in which the sediment slurries were housed and incubated in. These jars have a piece of cellulose filter paper affixed to the lid with epoxy, which was saturated with 1 ppt of zinc acetate solution in order to absorb any toxic hydrogen sulphide metabolite which was released during the incubation which may affect the results of the experiment. A strip of lead acetate paper was also affixed to the lid in order to visualise any $H_2S$ production. The structure of the jars used is shown in the Supplementary Figure S3.

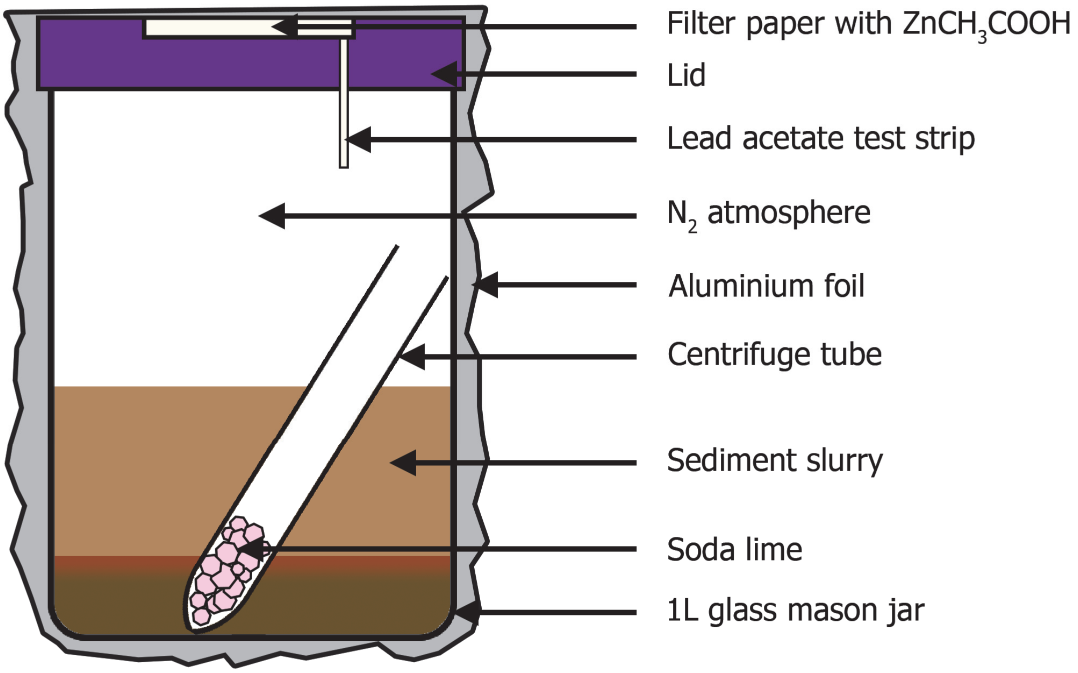


**Supplementary Figure S3:** The jars used in the incubation. The soda lime and zinc acetate impregnated filter paper serve to absorb toxic metabolites, while the N2 atmosphere within the jars ensures anoxic conditions are maintained.

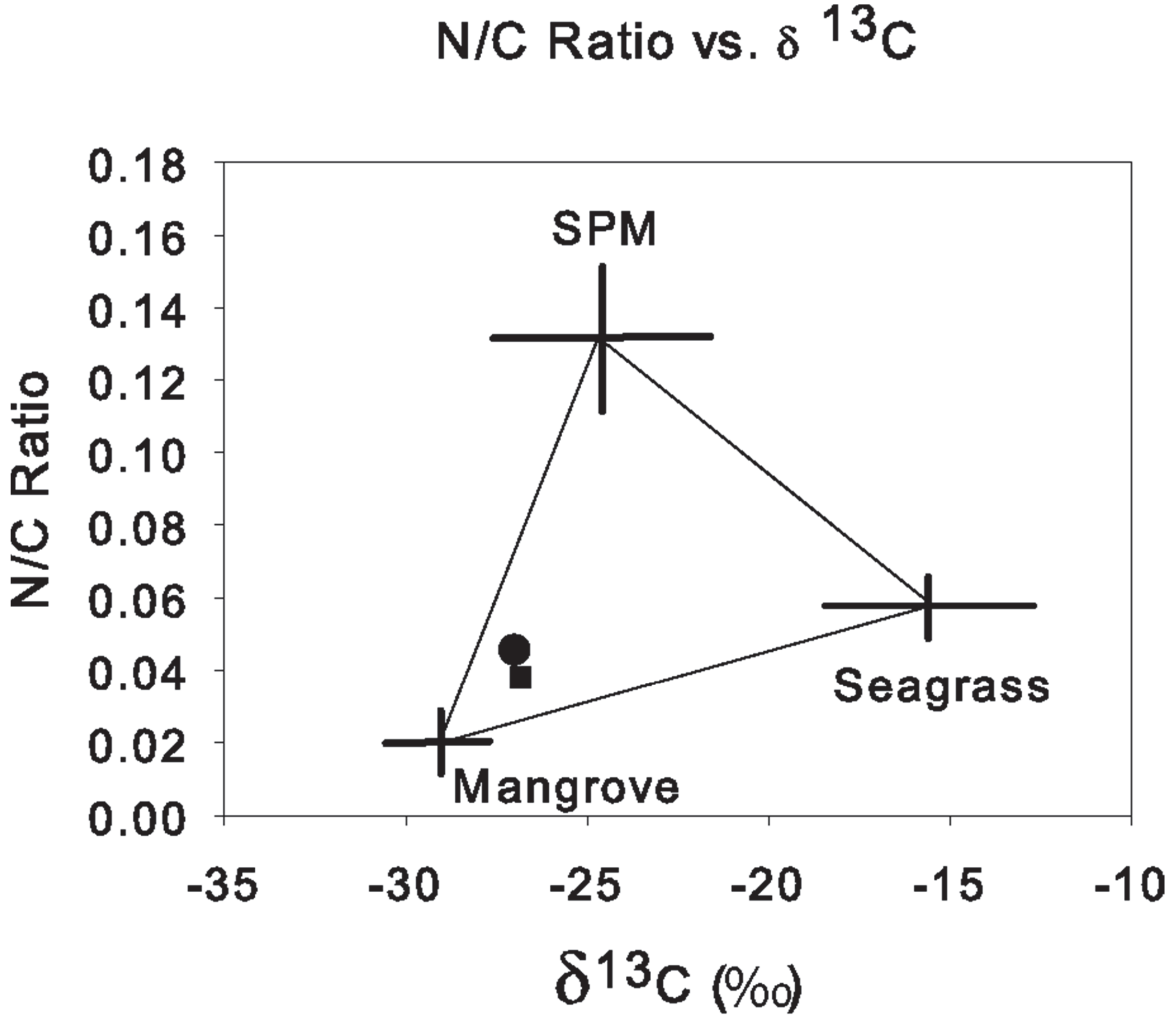


**Supplementary Figure S4:** N/C ratios of surface 2cm sediments, denoted by a circle, and sediments from the 20-22cm horizon, denoted by a square, against their 13C values. The crosses show the values of similar sources in a tropical Mexican mangrove (Eagle et al. 2004)

**Table S2:** POC content of the sediments throughout the anoxic incubation period and subsequent aeration incubation. S1, S2, S3 and S4 are the identifiers of the 4 replicates of the surface 2 cm. B1, B2, B3 and B4 are the identifiers of the 4 replicates of the sediments from 20-22 cm. All units are in mol $100g^{-1}$.

| Time (Days) | S1 | S2 | S3 | S4 | B1 | B2 | B3 | B4 |
|---|---|---|---|---|---|---|---|---|
| 0 | 0.6124 | 0.6124 | 0.5612 | 0.5684 | 0.5321 | 0.5321 | 0.5532 | 0.5532 |
| 7 | 0.5869 | 0.5614 | 0.5535 | 0.5691 | 0.5375 | 0.5391 | 0.5169 | 0.5182 |
| 21 | 0.5613 | 0.5646 | 0.5726 | 0.5626 | 0.5275 | 0.5433 | 0.5158 | 0.5389 |
| 42 | 0.5701 | 0.5756 | 0.5524 | 0.5683 | 0.5283 | 0.5270 | 0.5203 | 0.5465 |
| 63 | 0.5233 | 0.5246 | 0.5300 | 0.5456 | 0.4684 | 0.4562 | 0.4857 | 0.4854 |
| 105 | 0.5292 | 0.5246 | 0.5287 | 0.5230 | 0.5055 | 0.5003 | 0.4870 | 0.5228 |
| 140 | 0.5543 | 0.5569 | 0.5396 | 0.5204 | 0.4975 | 0.5070 | 0.4338 | 0.4698 |
| 175 | 0.4857 | 0.4878 | 0.4859 | 0.4946 | 0.4638 | 0.4727 | 0.4639 | 0.4666 |
| 210 | 0.4668 | 0.4640 | 0.4680 | 0.4745 | 0.4269 | 0.4220 | 0.4220 | 0.4364 |
| 308 | 0.4738 | -- | 0.4860 | 0.5025 | 0.4603 | 0.4629 | 0.4433 | 0.4619 |
| 365 | 0.4737 | -- | 0.5017 | 0.4653 | 0.4433 | 0.4513 | 0.4421 | 0.4467 |
| 400 | 0.4725 | -- | 0.4679 | 0.4655 | 0.4279 | 0.4381 | 0.4274 | 0.4185 |
| 470 | 0.4700 | -- | 0.4655 | 0.4592 | 0.4251 | 0.4587 | 0.4450 | 0.4509 |
| 500 | 0.4651 | -- | 0.4493 | 0.4688 | 0.4117 | 0.4021 | 0.4254 | 0.4152 |
| 531 | 0.4135 | -- | 0.3820 | 0.3706 | 0.3692 | 0.3346 | 0.3283 | 0.3892 |

**Table S3:** BOC content of the sediments throughout the anoxic incubation period and subsequent aeration incubation. S1, S2, S3 and S4 are the identifiers of the 4 replicates of the surface 2 cm. B1, B2, B3 and B4 are the identifiers of the 4 replicates of the sediments from 20-22 cm. All units are in mol $100g^{-1}$.

| Time (Days) | S1 | S2 | S3 | S4 | B1 | B2 | B3 | B4 |
|---|---|---|---|---|---|---|---|---|
| 0 | 0.0692 | 0.0692 | 0.0692 | 0.0692 | 0.0589 | 0.0589 | 0.0589 | 0.0589 |
| 7 | 0.0748 | 0.0750 | 0.0777 | 0.0848 | 0.0771 | 0.0770 | 0.0644 | 0.0729 |
| 21 | 0.0883 | 0.0849 | 0.0849 | 0.0939 | 0.0891 | 0.1052 | 0.0721 | 0.0882 |
| 42 | 0.1018 | 0.1126 | 0.1221 | 0.0986 | 0.0905 | 0.1122 | 0.0982 | 0.1010 |
| 63 | 0.0825 | 0.0909 | 0.0972 | 0.1084 | 0.0761 | 0.0762 | 0.0830 | 0.0931 |
| 105 | 0.0993 | 0.1118 | 0.1029 | 0.0982 | 0.1191 | 0.0909 | 0.1014 | 0.1042 |
| 140 | 0.1254 | 0.1302 | 0.1103 | 0.1055 | 0.0895 | 0.0910 | 0.0686 | 0.0523 |
| 175 | 0.0682 | 0.0637 | 0.0674 | 0.0693 | 0.0491 | 0.0701 | 0.0788 | 0.0529 |
| 210 | 0.0503 | 0.0485 | 0.0599 | 0.0467 | 0.0446 | 0.0428 | 0.0385 | 0.0358 |
| 308 | 0.0464 | -- | 0.0453 | 0.0496 | 0.0400 | 0.0393 | 0.0292 | 0.0407 |
| 365 | 0.0619 | -- | 0.0559 | 0.0578 | 0.0585 | 0.0551 | 0.0538 | 0.0613 |
| 400 | 0.0783 | -- | 0.0845 | 0.0898 | 0.0769 | 0.0839 | 0.0817 | 0.0899 |
| 470 | 0.0541 | -- | 0.0632 | 0.0578 | 0.0297 | 0.0334 | 0.0422 | 0.0415 |
| 500 | 0.0793 | -- | 0.0748 | 0.0719 | 0.0555 | 0.0048 | 0.0474 | 0.0465 |
| 531 | 0.1015 | -- | 0.0281 | 0.1049 | 0.0074 | 0.0260 | 0.0150 | 0.0322 |

**Table S4:** PIC content of the sediments throughout the anoxic incubation period and subsequent aeration incubation. S1, S2, S3 and S4 are the identifiers of the 4 replicates of the surface 2 cm. B1, B2, B3 and B4 are the identifiers of the 4 replicates of the sediments from 20-22 cm. All units are in mol 100g$^{-1}$.

| Time (Days) | S1 | S2 | S3 | S4 | B1 | B2 | B3 | B4 |
|---|---|---|---|---|---|---|---|---|
| 0 | 0.1117 | 0.1117 | 0.1117 | 0.1117 | 0.1121 | 0.1121 | 0.1121 | 0.1121 |
| 7 | 0.1389 | 0.1460 | 0.1438 | 0.1404 | 0.1371 | 0.1349 | 0.1364 | 0.146 |
| 21 | 0.1425 | 0.1426 | 0.1412 | 0.1397 | 0.1391 | 0.1335 | 0.1459 | 0.1376 |
| 42 | 0.1419 | 0.1373 | 0.1425 | 0.1608 | 0.1474 | 0.1417 | 0.1611 | 0.1377 |
| 63 | 0.1540 | 0.1533 | 0.1505 | 0.1436 | 0.1511 | 0.1444 | 0.1559 | 0.1507 |
| 105 | 0.1532 | 0.1483 | 0.1416 | 0.1501 | 0.1403 | 0.1395 | 0.1512 | 0.143 |
| 140 | 0.1437 | 0.1430 | 0.1730 | 0.1662 | 0.1230 | 0.1217 | 0.1616 | 0.1713 |
| 175 | 0.1272 | 0.1259 | 0.1302 | 0.1227 | 0.1291 | 0.1030 | 0.1252 | 0.1178 |
| 210 | 0.1307 | 0.1384 | 0.1303 | 0.1446 | 0.1302 | 0.1309 | 0.1292 | 0.1390 |
| 308 | 0.1376 | -- | 0.1334 | 0.1269 | 0.1466 | 0.1445 | 0.1425 | 0.1309 |
| 365 | 0.1292 | -- | 0.1820 | 0.2043 | 0.2063 | 0.2149 | 0.2104 | 0.2085 |
| 400 | 0.2164 | -- | 0.2152 | 0.2130 | 0.2310 | 0.228 | 0.2238 | 0.2171 |
| 470 | 0.2422 | -- | 0.2330 | 0.2362 | 0.2787 | 0.2392 | 0.2347 | 0.2416 |
| 500 | 0.2265 | -- | 0.2818 | 0.2418 | 0.2908 | 0.2841 | 0.2778 | 0.2899 |
| 531 | 0.1727 | -- | 0.1951 | 0.2507 | 0.1935 | 0.1669 | 0.1674 | 0.1520 |

**Table S5:** $CO_2$ absorbed by the soda lime placed within the headspace of the sediment slurry incubation bottles throughout the anoxic incubation period. S1, S3 and S4 are the identifiers of the 3 replicates of the surface 2 cm. B1, B2, B3 and B4 are the identifiers of the 4 replicates of the sediments from 20-22 cm. All units are in mol 100g dry weight of sediment$^{-1}$.

| Time (Days) | S1 | S3 | S4 | B1 | B2 | B3 | B4 |
|---|---|---|---|---|---|---|---|
| 0 | 0.0000 | 0.0000 | 0.0000 | 0.0000 | 0.0000 | 0.0000 | 0.0000 |
| 7 | 0.0170 | 0.0115 | 0.0104 | 0.0074 | 0.0071 | 0.0067 | 0.0068 |
| 21 | 0.0345 | 0.0277 | 0.0278 | 0.0173 | 0.0170 | 0.0165 | 0.0177 |
| 42 | 0.0544 | 0.0471 | 0.0498 | 0.0310 | 0.0307 | 0.0307 | 0.0330 |
| 63 | 0.0753 | 0.0667 | 0.0658 | 0.0436 | 0.0421 | 0.0427 | 0.0444 |
| 105 | 0.1032 | 0.1004 | 0.0964 | 0.0682 | 0.0654 | 0.0674 | 0.0689 |
| 140 | 0.1343 | 0.1349 | 0.1252 | 0.0887 | 0.0874 | 0.0901 | 0.1704 |
| 175 | 0.1599 | 0.1658 | 0.1531 | 0.1115 | 0.3222 | 0.2182 | 0.1951 |
| 210 | 0.1836 | 0.1903 | 0.1551 | 0.1303 | 0.3141 | 0.2359 | 0.1944 |
| 308 | 0.2347 | 0.2509 | 0.2569 | 0.1677 | 0.3819 | 0.2805 | 0.2707 |
| 365 | 0.2804 | 0.2464 | 0.2170 | 0.1427 | 0.3414 | 0.5471 | 0.2308 |
| 400 | 0.2575 | 0.2971 | 0.2645 | 0.1794 | 0.3541 | 0.5869 | 0.3038 |
| 470 | 0.2817 | 0.3271 | 0.2924 | 0.1783 | 0.3906 | 0.6026 | 0.3266 |
| 500 | 0.3246 | 0.3029 | 0.3133 | 0.1907 | 0.3704 | 0.5842 | 0.3026 |

**Table S6:** Ammonia content of the porewater of the sediment slurry throughout the anoxic incubation period and subsequent aeration incubation. S1, S2, S3 and S4 are the identifiers of the 4 replicates of the surface 2 cm. B1, B2, B3 and B4 are the identifiers of the 4 replicates of the sediments from 20-22 cm. All units are in mmol 100g dry weight of sediment$^{-1}$.

| Time (Days) | S1 | S2 | S3 | S4 | B1 | B2 | B3 | B4 |
|---|---|---|---|---|---|---|---|---|
| 0 | 0.0877 | 0.0877 | 0.0877 | 0.0877 | 0.1391 | 0.1391 | 0.1391 | 0.1391 |
| 7 | 0.0970 | 0.0939 | 0.0927 | 0.1038 | 0.0749 | 0.0636 | 0.0647 | 0.0740 |
| 21 | 0.1066 | 0.1031 | 0.1019 | 0.1141 | 0.0821 | 0.0697 | 0.0709 | 0.0812 |
| 42 | 0.0208 | 0.0332 | 0.0290 | 0.0667 | 0.0093 | 0.0061 | 0.0072 | 0.0069 |
| 63 | 0.0223 | 0.0220 | 0.0207 | 0.0274 | 0.0090 | 0.0072 | 0.0066 | 0.0113 |
| 105 | 0.0960 | 0.0371 | 0.0772 | 0.0316 | 0.0138 | 0.0105 | 0.0176 | 0.0101 |
| 140 | 0.0215 | 0.0707 | 0.0231 | 0.0327 | 0.0092 | 0.0128 | - | 0.0081 |
| 175 | 0.0565 | 0.0404 | 0.0207 | 0.0262 | 0.0091 | 0.0095 | 0.0117 | 0.0120 |
| 210 | 0.0627 | 0.0680 | 0.0663 | 0.0669 | 0.0099 | 0.0092 | 0.0066 | 0.0069 |
| 308 | 0.1958 | - | 0.2479 | 0.1682 | 0.0711 | 0.1130 | 0.0874 | 0.0813 |
| 365 | 0.0607 | - | 0.0746 | 0.0749 | 0.0228 | 0.0165 | 0.0178 | 0.0123 |
| 400 | 0.0924 | - | 0.1184 | 0.1073 | 0.0277 | 0.0199 | 0.0352 | 0.0229 |
| 470 | 0.1340 | - | 0.0966 | 0.1268 | 0.0454 | 0.0576 | 0.0700 | 0.0490 |
| 500 | 0.1657 | - | 0.1925 | 0.1782 | 0.0710 | 0.0689 | 0.0786 | 0.0431 |
| 531 | 0.0249 | - | 0.0429 | 0.0578 | 0.0579 | 0.0317 | 0.0092 | 0.0254 |

**Table S7:** The 355nm, 400nm, 440 nm and 550nm absorbance values of the porewater of the surface sediment slurry throughout the anoxic incubation period and subsequent aeration incubation. S1, S2, S3 and S4 are the identifiers of the 4 replicates of the surface 2 cm. Blank spaces represent samples which did not provide a reliable reading.

| Time | 355 nm absorbance | | | | 400 nm absorbance | | | | 440 nm absorbance | | | | 550 nm absorbance | | | |
|---|---|---|---|---|---|---|---|---|---|---|---|---|---|---|---|---|
| (days) | S1 | S2 | S3 | S4 | S1 | S2 | S3 | S4 | S1 | S2 | S3 | S4 | S1 | S2 | S3 | S4 |
| 0 | 0.057 | 0.057 | 0.057 | 0.057 | 0.039 | 0.039 | 0.039 | 0.039 | 0.022 | 0.022 | 0.022 | 0.022 | 0.008 | 0.008 | 0.008 | 0.008 |
| 7 | 0.068 | 0.082 | -- | 0.071 | 0.050 | 0.060 | -- | 0.051 | 0.029 | 0.035 | -- | 0.028 | 0.013 | 0.016 | -- | 0.011 |
| 21 | 0.070 | 0.075 | -- | 0.072 | 0.049 | 0.051 | -- | 0.052 | 0.027 | 0.028 | -- | 0.030 | 0.011 | 0.011 | -- | 0.013 |
| 42 | -- | 0.087 | 0.082 | 0.081 | 0.048 | 0.043 | 0.055 | 0.058 | -- | 0.038 | 0.030 | 0.035 | -- | 0.018 | 0.011 | 0.017 |
| 63 | 0.084 | 0.063 | 0.059 | 0.072 | -- | 0.061 | 0.045 | 0.056 | 0.028 | 0.033 | 0.035 | 0.033 | 0.015 | 0.015 | 0.009 | 0.011 |
| 105 | 0.057 | 0.060 | 0.033 | 0.076 | 0.064 | 0.049 | 0.036 | 0.048 | -- | 0.017 | 0.035 | 0.023 | -- | 0.006 | 0.007 | 0.026 |
| 140 | 0.057 | 0.058 | -- | 0.044 | 0.036 | 0.033 | -- | 0.027 | 0.027 | 0.022 | 0.039 | 0.017 | 0.015 | 0.024 | 0.012 | 0.008 |
| 175 | 0.067 | 0.055 | 0.065 | 0.062 | 0.047 | 0.034 | 0.042 | 0.039 | 0.036 | 0.023 | 0.028 | 0.028 | 0.022 | 0.014 | 0.015 | 0.016 |
| 210 | 0.077 | 0.073 | 0.080 | 0.084 | 0.051 | 0.059 | 0.057 | 0.069 | 0.040 | 0.037 | 0.037 | 0.033 | 0.025 | 0.017 | 0.017 | 0.022 |
| 308 | 0.082 | -- | 0.084 | 0.076 | 0.048 | -- | 0.046 | 0.039 | 0.032 | -- | 0.029 | 0.025 | 0.014 | -- | 0.011 | 0.013 |
| 365 | 0.085 | -- | 0.057 | 0.065 | 0.051 | -- | 0.029 | 0.034 | 0.037 | -- | 0.018 | 0.023 | 0.021 | -- | 0.008 | 0.011 |
| 400 | 0.069 | -- | 0.071 | 0.073 | 0.049 | -- | 0.037 | 0.033 | 0.008 | -- | 0.018 | 0.018 | 0.012 | -- | 0.008 | 0.007 |
| 470 | 0.062 | -- | 0.063 | 0.073 | 0.031 | -- | 0.029 | 0.033 | 0.010 | -- | 0.010 | 0.019 | 0.007 | -- | 0.005 | 0.011 |
| 500 | 0.080 | -- | 0.074 | 0.076 | 0.041 | -- | 0.035 | 0.039 | 0.007 | -- | 0.009 | 0.007 | 0.007 | -- | 0.005 | 0.011 |
| 531 | 0.104 | -- | 0.076 | 0.134 | 0.078 | -- | 0.056 | 0.099 | 0.569 | -- | 0.726 | 0.840 | 0.034 | -- | 0.025 | 0.033 |

**Table S8:** The 355nm, 400nm, 440 nm and 550nm absorbance values of the porewater of the surface sediment slurry throughout the anoxic incubation period and subsequent aeration incubation. B1, B2, B3 and B4 are the identifiers of the 4 replicates of the sediments from 20-22 cm. Blank spaces represent samples which did not provide a reliable reading.

| Time | 355 nm absorbance | | | | 400 nm absorbance | | | | 440 nm absorbance | | | | 550 nm absorbance | | | |
|---|---|---|---|---|---|---|---|---|---|---|---|---|---|---|---|---|
| (days) | B1 | B2 | B3 | B4 | B1 | B2 | B3 | B4 | B1 | B2 | B3 | B4 | B1 | B2 | B3 | B4 |
| 0 | 0.052 | 0.052 | 0.052 | 0.052 | 0.031 | 0.031 | 0.031 | 0.031 | 0.021 | 0.021 | 0.021 | 0.021 | 0.012 | 0.012 | 0.012 | 0.012 |
| 7 | -- | 0.071 | 0.076 | 0.067 | -- | 0.049 | 0.052 | 0.043 | -- | 0.036 | 0.036 | 0.029 | -- | 0.020 | 0.019 | 0.015 |
| 21 | 0.044 | -- | 0.064 | 0.068 | 0.026 | -- | 0.045 | 0.062 | 0.015 | -- | 0.032 | 0.026 | 0.006 | -- | 0.019 | 0.019 |
| 42 | 0.070 | -- | 0.074 | 0.083 | 0.048 | -- | 0.051 | 0.059 | 0.034 | -- | 0.036 | 0.043 | 0.009 | -- | 0.009 | 0.015 |
| 63 | -- | 0.084 | -- | 0.081 | -- | 0.064 | -- | 0.059 | -- | 0.036 | -- | 0.029 | -- | 0.020 | -- | 0.015 |
| 105 | 0.068 | 0.083 | 0.074 | 0.072 | 0.049 | 0.052 | 0.055 | 0.053 | 0.039 | 0.049 | 0.045 | 0.042 | 0.018 | 0.019 | 0.021 | 0.020 |
| 140 | 0.078 | 0.062 | -- | 0.064 | 0.058 | 0.042 | -- | 0.045 | 0.047 | 0.031 | -- | 0.036 | 0.024 | 0.019 | -- | 0.024 |
| 175 | 0.079 | 0.043 | 0.082 | 0.075 | 0.059 | 0.025 | 0.058 | 0.043 | 0.048 | 0.017 | 0.047 | 0.045 | 0.025 | 0.010 | 0.023 | 0.019 |
| 210 | 0.069 | 0.073 | 0.052 | 0.061 | 0.054 | 0.059 | 0.064 | 0.046 | 0.039 | 0.041 | 0.033 | 0.032 | 0.016 | 0.028 | 0.016 | 0.009 |
| 308 | 0.074 | 0.06 | 0.072 | 0.064 | 0.043 | 0.031 | 0.039 | 0.036 | 0.029 | 0.02 | 0.026 | 0.025 | 0.014 | 0.01 | 0.012 | 0.013 |
| 365 | 0.084 | 0.083 | 0.061 | 0.048 | 0.051 | 0.038 | 0.029 | 0.037 | 0.042 | 0.024 | 0.016 | 0.014 | 0.028 | 0.017 | 0.009 | 0.007 |
| 400 | 0.087 | 0.072 | 0.075 | 0.087 | 0.048 | 0.036 | 0.049 | 0.035 | 0.019 | 0.017 | 0.021 | 0.019 | 0.014 | 0.012 | 0.012 | 0.008 |
| 470 | 0.042 | 0.052 | 0.047 | 0.041 | 0.029 | 0.024 | 0.021 | 0.016 | 0.012 | 0.015 | 0.01 | 0.007 | 0.011 | 0.009 | 0.005 | 0.003 |
| 500 | 0.082 | 0.083 | 0.084 | 0.080 | 0.033 | 0.035 | 0.029 | 0.049 | 0.014 | 0.013 | 0.015 | 0.013 | 0.009 | 0.007 | 0.007 | 0.008 |
| 531 | 0.104 | 0.166 | 0.162 | 0.125 | 0.092 | 0.138 | 0.098 | 1.562 | 0.501 | 0.845 | 1.419 | 0.387 | 0.066 | 0.107 | 0.043 | 0.021 |

**Table S9:** pH of the porewater of the sediment slurry throughout the anoxic incubation period and subsequent aeration incubation. S1, S2, S3 and S4 are the identifiers of the 4 replicates of the surface 2 cm. B1, B2, B3 and B4 are the identifiers of the 4 replicates of the sediments from 20-22 cm.

| Time (Days) | S1 | S2 | S3 | S4 | B1 | B2 | B3 | B4 |
|---|---|---|---|---|---|---|---|---|
| 105 | 7.4 | 7.4 | 7.7 | 7.1 | 6.0 | 5.7 | 5.6 | 5.5 |
| 140 | 7.0 | 7.4 | 7.3 | 7.1 | 6.5 | 5.8 | 5.7 | 5.8 |
| 175 | 6.7 | 7.1 | 6.9 | 6.1 | 5.7 | 6.3 | 6.2 | 5.9 |
| 210 | 7.3 | 6.9 | 7.3 | 7.1 | 6.2 | 5.9 | 6.0 | 5.8 |
| 308 | 5.2 | -- | 5.3 | 5.6 | 5.5 | 5.7 | 5.8 | 5.2 |
| 365 | 5.3 | -- | 5.2 | 5.1 | 5.2 | 5.5 | 4.9 | 5.3 |
| 400 | 5.5 | -- | 5.7 | 5.1 | 5.1 | 5.1 | 5.6 | 5.2 |
| 470 | 5.6 | -- | 5.3 | 5.7 | 5.4 | 5.4 | 5.0 | 5.8 |
| 500 | 5.6 | -- | 5.4 | 5.7 | 5.3 | 5.5 | 5.3 | 6.0 |

**Table S10:** Iron content of the sediments throughout the anoxic incubation period and subsequent aeration incubation. S1, S3 and S4 are the identifiers of 3 replicates of the surface 2 cm. B1, B2, and B3 are the identifiers of 3 replicates of the sediments from 20-22 cm. All units are in mol $100g^{-1}$.

| Time (days) | S1 | S3 | S4 | B1 | B2 | B3 |
|---|---|---|---|---|---|---|
| 7 | 0.0542 | 0.0562 | 0.0522 | 0.0448 | 0.0448 | 0.0410 |
| 21 | 0.0591 | 0.0512 | 0.0521 | 0.0504 | 0.0504 | 0.0455 |
| 42 | 0.0476 | 0.0538 | 0.0523 | 0.0532 | 0.0532 | 0.0436 |
| 63 | 0.0537 | 0.0554 | 0.0537 | 0.0427 | 0.0427 | 0.0502 |
| 147 | 0.0535 | 0.0554 | 0.0535 | 0.0457 | 0.0457 | 0.0443 |
| 231 | 0.0533 | 0.0571 | 0.0528 | 0.0457 | 0.0457 | 0.0458 |
| 329 | -- | 0.0513 | 0.0560 | 0.0433 | 0.0433 | 0.0523 |
| 365 | 0.0565 | 0.0551 | 0.0528 | 0.0472 | 0.0472 | 0.0537 |
| 400 | 0.0575 | 0.0556 | 0.0526 | 0.0468 | 0.0468 | 0.0482 |
| 500 | 0.0803 | 0.0558 | 0.0575 | 0.0494 | 0.0494 | 0.0465 |

**Table S11:** Uncorrected stable isotopes data of $^{13}C$ and $^{15}N$ of selected sediments used in the incubation experiment. S 0 and B 0 are the samples of the surface 2 cm and sediments from 20-22 cm at the start of the incubation, while S1 210, S2 210, S3 210 and S4 210 are the identifiers of the 4 replicates of the surface 2 cm after 210 days of anoxic incubation. B1 210, B2 210, B3 210 and B4 210 are the identifiers of the 4 replicates of the sediments from 20-22 cm after 210 days of anoxic incubation.

| Sample | $\delta^{13}C$ (‰) | $\delta^{15}N$ (‰) | %C | %N | N/C Ratio |
|---|---|---|---|---|---|
| S 0 | -26.92 | 4.18 | 7.67 | 0.37 | 0.066 |
| B 0 | -26.70 | 2.21 | 7.25 | 0.28 | 0.071 |
| S1 210 | -27.00 | 4.33 | 7.30 | 0.33 | 0.066 |
| S1 210 | -26.99 | 4.05 | 7.44 | 0.34 | 0.065 |
| S2 210 | -27.02 | 4.01 | 7.67 | 0.35 | 0.064 |
| S3 210 | -27.00 | 4.02 | 7.48 | 0.34 | 0.065 |
| S4 210 | -27.08 | 4.08 | 7.53 | 0.34 | 0.064 |
| B1 210 | -26.86 | 2.40 | 7.35 | 0.28 | 0.071 |
| B2 210 | -26.90 | 2.26 | 7.30 | 0.28 | 0.071 |
| B3 210 | -26.85 | 2.31 | 7.55 | 0.28 | 0.069 |
| B4 210 | -26.84 | 2.45 | 7.33 | 0.28 | 0.071 |
